# Geometric control of Myosin-II orientation during axis elongation

**DOI:** 10.1101/2022.01.12.476069

**Authors:** M.F. Lefebvre, N.H. Claussen, N.P. Mitchell, H.J. Gustafson, S.J. Streichan

## Abstract

The actomyosin cytoskeleton is a crucial driver of morphogenesis. Yet how the behavior of large-scale cytoskeletal patterns in deforming tissues emerges from the interplay of geometry, genetics, and mechanics remains incompletely understood. Convergent extension flow in *D. melanogaster* embryos provides the opportunity to establish a quantitative understanding of the dynamics of anisotropic non-muscle myosin II. Cell-scale analysis of protein localization in fixed embryos suggests that there are complex rules governing how the control of myosin anisotropy is regulated by gene expression patterns. However, technical limitations have impeded quantitative and dynamic studies of this process at the whole embryo level, leaving the role of geometry open. Here we combine *in toto* live imaging with quantitative analysis of molecular dynamics to characterize the distribution of myosin anisotropy and corresponding genetic patterning. We found pair rule gene expression continuously deformed, flowing with the tissue frame. In contrast, myosin anisotropy orientation remained nearly static, aligned with the stationary dorsal-ventral axis of the embryo. We propose myosin recruitment by a geometrically defined static source, potentially related to the embryo-scale epithelial tension, and account for transient deflections by the interplay of cytoskeletal turnover with junction reorientation by flow. With only one parameter, this model quantitatively accounts for the time course of myosin anisotropy orientation in wild-type, twist, and even-skipped embryos as well as embryos with perturbed egg geometry. Geometric patterning of the cytoskeleton suggests a simple physical strategy to ensure a robust flow and formation of shape.

## INTRODUCTION

During morphogenesis, tissues dynamically remodel through cellular flows [1]. These flows are driven by patterned cytoskeletal processes, such as large-scale gradients of non-muscle myosin II (myosin), that generate imbalanced forces [2]. Two processes affect gene expression and cytoskeletal patterns during morphogenesis. First, cells move, taking their constituents with them. Second, the contents of cells are constantly reorganizing [3, 4]. If intracellular turnover is slow compared to the rate of tissue movement, pattern change is dominated by advection. In fluid dynamics, such patterns are referred to as “Lagrangian” [5]. Recent technological advances now allow the study of dynamic patterns at a global scale. In this way, it becomes possible to elucidate the effects of turnover and tissue-scale cues relating dynamics of gene expression patterns, cytoskeletal components, and tissue shape during morphogenesis.

Here we analyze the dynamics of myosin in *D. melanogaster* to establish a quantitatively testable link between genetic patterning and organ geometry. We study the dynamics of the anisotropic distribution of myosin which drives global tissue flow through cell intercalation during body axis elongation [6–9], known as germband extension (GBE). Qualitative analysis of fixed embryos has been used to suggest a link between spatial patterns of pair rule genes (PRGs) and the orientation of myosin rich junctions (MRJs), likely via members of the Toll-like transmembrane receptor (TLRs) family [10–14]. However, cell-scale live imaging demonstrates reorientation of myosin anisotropy by other factors, for example mechanosensation (Fig. 1a) [15, 16].

**FIG. 1.**
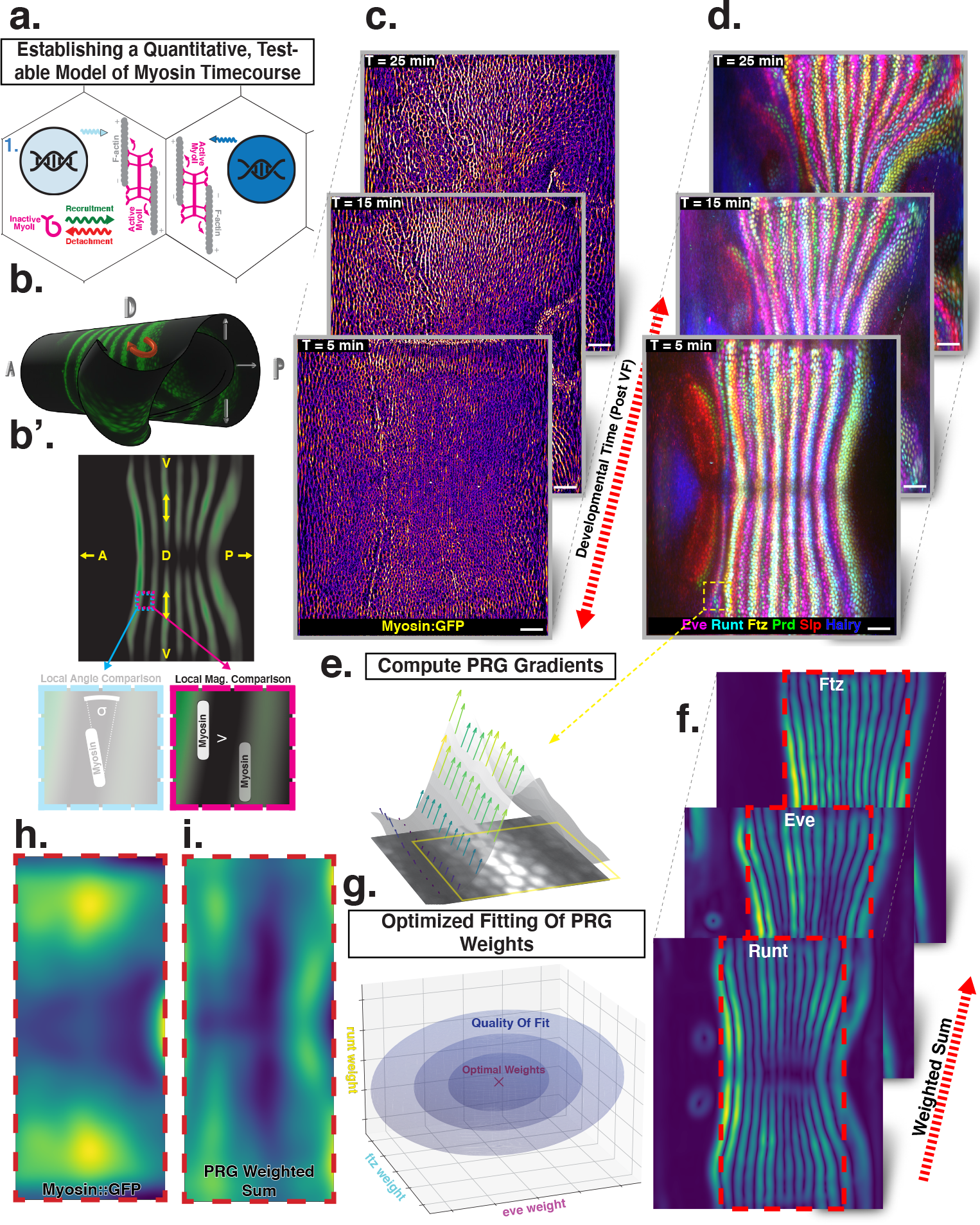
Global analysis of myosin vs PRG expression patterns reveals no linear correlation. (a) Junctional myosin could be regulated by gene expression patterns, by mechanical cues, or both. (b-b’) Tissue cartography extracts embryo surfaces from volumetric *in toto* light sheet imaging which are projected onto a cylindrical chart. This allows measuring quantities on a tissue scale (b), here the intensity and orientation of junctional myosin and PRGs. (c) Time series of junctional myosin, starting at 5 minutes post ventral furrow initiation (d) Time series of PRGs, starting at 5 minutes post ventral furrow initiation. Time series created by digitally stitching together different stained and live imaged embryos. (e) The smoothed gradient of PRG expression patterns computes local cell-cell differences. The gradient vector points in direction in which the signal increases most. (f) Gradient magnitude of the expression patterns of the PRGs Runt, Eve, and Ftz in 3 representative embryos shows 14 stripes with a DV modulation of intensity. (g) In regression analysis, the smoothed gradients of PRGs are combined into a weighted sum to approximate the observed myosin pattern. The weights are adjusted to optimize the quality of fit, exploring the entire space of possible weights, both positive and negative. (h) Smoothed junctional myosin intensity at the onset of GBE (ensemble average of 5 embryos). (i) Result of PRG gradient regression. The best possible fit using the weighted sum of PRG gradients does not resemble the large-scale myosin pattern.

Using *in toto* light sheet microscopy and tissue cartography [17, 18], we map the time course of myosin, PRGs, and TLRs during GBE (Fig. 1c-d). These maps show quantitatively that PRG and TLR expression patterns deform with tissue flow, whereas myosin orientation is only transiently deflected away from stationary geometric landmarks in response to flow. This leads to an increasing discrepancy between the pattern of PRG expression and myosin orientation over the course of GBE. We quantitatively explain the short-lived anisotropy deflection by the finite time of association (∼ 5 minutes) between myosin motors remain and the actin cortex. These results demonstrate that PRGs and TLRs are in a flowing frame of reference (Lagrangian), while the recruitment of myosin driving flow is controlled by a nearly static frame.

## RESULTS

### A quantitative mismatch between junctional myosin accumulation and PRG gradient patterns

Despite extensive analysis [11–13, 19], little is known about the quantitative dynamics of both myosin and PRGs at the whole embryo level during GBE. We digitally stitched data gathered from multiple live and fixed embryos to create a dynamic atlas comprising components of the anterior-posterior (AP) patterning system as well as myosin (Fig. 1 c-d), measured across the entire embryo [20]. The time course of these gene products is provided with ∼1 minute temporal resolution, starting from cellularization until the end of GBE (See SI for detail). We define t = 0 to be the initiation of ventral furrow (VF) formation.

Throughout embryogenesis, egg geometry remains static (Fig. 1b), defining a fixed reference frame that is described by a coordinate system parallel to the anterior-posterior (AP) and dorsal-ventral (DV) axes. We focus on junctional myosin at the apical surface [9] (Fig. 1c), together with the PRGs Even-Skipped (Eve), Runt, Fushi Tarazu (Ftz), Hairy, Paired (Prd), and Sloppy-Paired (Slp) to create a dynamic atlas of gene expression during GBE (Fig. 1d). As seen previously, MRJs mainly align with the DV axis (Fig. 1c) [6], while all of the PRGs we analyzed were expressed in a series of stripes occurring at regular intervals along the AP axis (Fig. 1d) [10, 21]. We mine this expression atlas to quantitatively test the relationship between the accumulation of myosin on junctions and cumulative PRG expression (see SI Sect. I A 4).

Differences of PRG expression levels in adjacent cells have been proposed to underlie anisotropic myosin accumulation [9, 11]. This provides a testable prediction relating PRGs to the accumulated signal and orientation of MRJs. We first computed local differences (gradients) of PRG expression levels, focusing on Runt and Eve – which are known to have the strongest impact on GBE [10] – as well as Ftz. Gradients were steepest along the AP axis, pointing towards the center of the stripes (Fig. 1e). Across the germband, the magnitude of differences showed 14 regularly-spaced stripes. While all PRGs are slightly out of register with one another [21], the six PRGs we analyzed had the following characteristics: (i) along the AP axis, gradients were strongest at the first and last stripe [22], (ii) expression levels reduced towards the dorsal pole (Fig. 1f).

The profile of junctional myosin also demonstrated a DV gradient similar to the one observed for the PRGs, with a minimum on the dorsal pole [23]. However, along the AP axis, the myosin profile did not match any individual PRG gradient, because they all have gaps in intensity between stripes (Fig. 1c), while myosin does not. Therefore, we investigated whether combining the gradient profiles of multiple PRGs could produce a pattern consistent with the observed myosin accumulation. We used linear regression to compare the observed magnitude of junctional myosin with a weighted sum of PRG gradient patterns. The weighting parameters are adjusted to achieve the best possible agreement with the myosin profile (Eq. 1).

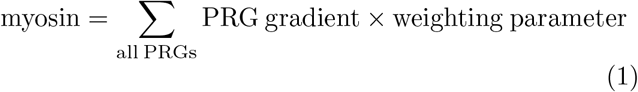

Each PRG had its own parameter, which does not change over space or between different stripes. It can be positive or negative, representing a promotion or inhibition of myosin accumulation (for details, see SI Sect. I D). The PRG regression can also be seen as an analysis of the large-scale correlation between junctional myosin and PRG gradients, without making any *a priori* assumptions about the individual effects of each PRG.

The best fit produced in this way captured the observed DV modulation of myosin (Figs. 1h,i). Along the AP axis, however, the patterns were quantitatively and qualitatively different. The model of local PRG differences predicted strong myosin accumulations at the anterior and posterior ends of the germband and a minimum in the center (Fig. 1i). This did not fit the observed myosin localization pattern (Fig. 1h). Our analysis suggests that local differences of PRGs cannot be linearly related to the amount of myosin accumulating on junctions, without postulating currently unknown additional complexity.

### Reorientation of PRGs by advection vs. near-stationary orientation of myosin anisotropy

Anisotropic myosin distributions are characterized by both a local intensity – which did not correlate with PRG patterns – and a local orientation, to which we turn next. In any version of instructive genetic regulation of myosin recruitment, the angle between the myosin orientation and the gene stripes ought to be constant over all times. Therefore, we analyzed the local orientation of MRJs across the whole embryo surface over time, which we compared to the concurrent orientation of local PRG stripes.

During GBE, both MRJs and PRG orientations are under the influence of significant tissue flow. As a consequence, their patterns could deform with the flow (i.e Lagrangian behavior). We characterized the time course of the instantaneous flow field, which quantifies the global pattern of cell movements. We compared this quantification of flow with the temporal evolution localization patterns of PRGs and TLR (Figs. 2 a-c’).

**FIG. 2.**
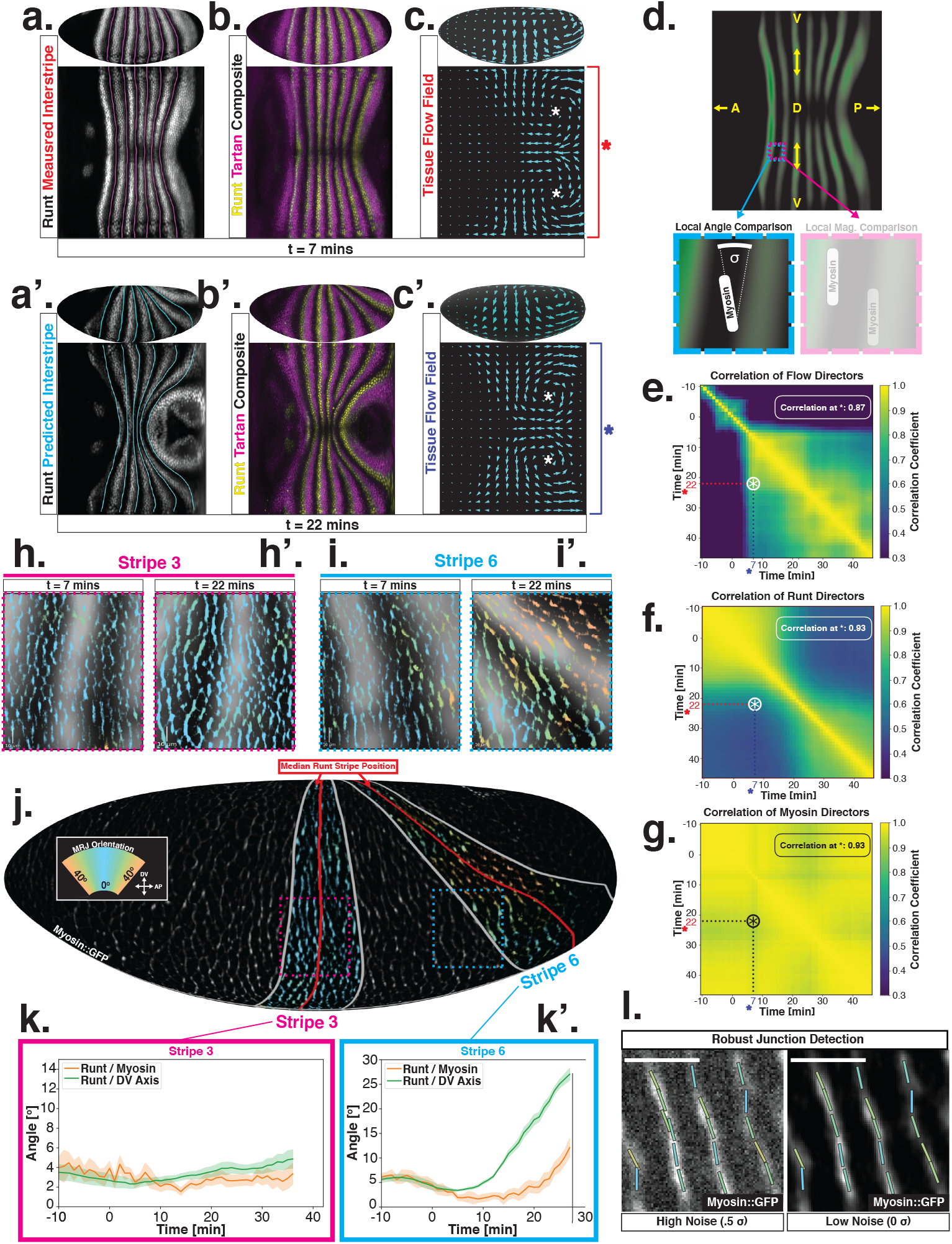
PRGs flow with tissue while myosin pattern does not. (a) Runt stripes with measured inter-stripe lines, 7 minutes post VF initiation from a representative Runt::LlamaTag-GFP embryo. All PRG stripes are initially approximately parallel to the DV axis. (a’) PRG stripes deform due to advection by tissue flow. Runt stripes with inter-stripe lines predicted by advection, 22 minutes post VF initiation. Same embryo as in (A). (b-b’) Digitally stitched Runt/Tartan composite, 7 minutes (b) and 22 minutes (b’) post VF initiation, showing that PRG and TLR stripes remain parallel. (c-c’) Tissue flow field, 7 minutes (c) and 22 minutes (c’) post VF initiation. Calculated from an average of 5 WT Myosin::GFP embryos. (d) Using tissue cartography, we compare MRJ and PRG orientation across the entire embryo. (e) Temporal autocorrelation of the tissue flow field. Each pixel in the matrix shows the correlation (similarity in direction, ranging from 0 to 1, averaged over the embryo surface) of the flow fields at two different time points. (f) Temporal autocorrelation of the Runt stripe direction shows rapid decay during tissue flow. Data from 5 WT Runt::LlamaTag-GFP embryos (see SI Sect I A 5 and I A 4 c for mathematical details). (g) Temporal autocorrelation of the tissue-scale myosin direction, showing an approximately static pattern of myosin orientation. Data from 5 WT Myosin::GFP embryos. (h’-h’, i-i’) Digitally stitched images showing Runt and junctional myosin (colored according to angle with DV axis) in a part of the regions surrounding runt stripes 3 (h-h’) and 6 (i-i’) at 7 minutes (h,i) and 22 minutes (h’,i’) post VF initiation. In regions were gene patterns are deformed by flow, an angle discrepancy between myosin orientation and PRG stripes develops. (j) Junctional myosin at 22 minutes post VF initiation in a representative WT Myosin::GFP embryo. In highlighted regions (defined by Runt stripes 3 and 6), junction color corresponds to the junction/DV axis angle and junction brightness to the myosin fluorescent intensity. Red lines show median runt stripe position. (k-k’) Angle between myosin anisotropy orientation and Runt stripe, and angle between Runt stripe and DV axis, averaged over the regions corresponding to runt stripes 3 and 6. Runt angle measured by the direction of Runt gradient, rotated by 90°. (l) The radon transform method detects MRJs and is insensitive to noise. Bars indicate junctions detected by the radon transform, color-coded according to their angle.

Just after the initiation of gastrulation (7 minutes post VF initiation), Runt localization in the germband is characterized by seven stripes at different AP positions (Figs. 2a,b, SI) [21]. Since this pattern of expression is stereotypic for the PRGs known to have the largest individual effects on GBE [10, 21] we adopted Runt as our representative PRG. 15 minutes later, (22 minutes post VF initiation), the PRG and TLR stripes were strongly deformed, with the posterior-most Runt stripe (stripe seven) almost completely moved onto the dorsal side of the embryo. We also tested whether the dynamic expression of TLRs matched what is observed for PRGs. We found that the orientation of Tartan stripes [13], closely mirrored that of the Runt stripes (Figs. 2a’,b’, SI).

The reorientation of PRG and TLR localization during GBE is reflected in a rapid decline of the autocorrelation of local stripe orientations (Fig. 2f). To understand whether tissue flow could account for this deformation, we calculated six inter-stripe positions of the Runt pattern (Fig. 2a, magenta lines, SI Fig. 8) which we advected along cell trajectories computed from the velocity field, as measured by particle image velocimetry (PIV). The resulting advected inter-stripe positions (Fig. 2a’, cyan lines) remained located between the position of Runt stripes at 22 minutes. Manual tracking (SI Fig. 9) confirms that cells which initially express Runt retain expression after 20 minutes. Together, this suggests that both PRG and TLR patterns flow with the tissue frame of reference and are thus Lagrangian: their reorientation is quantitatively accounted for by advection due to tissue flow.

Next, we asked whether the velocity field behaved in the same manner. By 7 minutes post VF initiation, a characteristic tissue flow pattern emerged, with four vortices and two hyperbolic fixed points [23] (Fig. 2c). This characteristic flow pattern persisted during GBE, as highlighted by a very similar pattern that is observed 15 minutes later (Fig. 2c’). While the magnitude of the flow clearly changed, its local direction was nearly constant, and vortices shifted only slightly over time. The auto-correlation of local tissue flow directions remained high throughout GBE (Fig. 2e). Thus, the instantaneous flow field was nearly stationary for the entire duration of GBE: although cells travel long distances across the embryo surface, different cell neighborhoods that pass through a common spatial coordinate will move in the same direction, irrespective of the precise time point during GBE.

Together, these results raise a problem. Tissue flow is known to be driven by anisotropic myosin [6–9]. In fact, the flow field can be quantitatively predicted from the myosin distribution using a simple hydrodynamic model [23]. This suggests that a static myosin pattern should be required to produce the observed, nearly static, tissue flow field. Yet directional myosin recruitment by cell-intrinsic PRG patterning should lead to a continuous reorientation and advection of myosin anisotropy.

To resolve this discrepancy, we measured the orientation of MRJs and compared them to local Runt stripe orientations (Figs. 2h-k’). The latter is defined by the direction of the Runt gradient, rotated by 90°. MRJs were detected using a segmentation free method based on the Radon transform (Figs. 2j,l) [23] (see SI Sect. I A 4). This method is robust even at low signal-to-noise ratios. The local myosin anisotropy direction (Figs 2k-k’) was defined by the intensity-weighted mean of MRJ orientations in a 3-cell radius (see SI Sect. I A 4). To link local angles with the organ-scale geometry, we measured angles with respect to the DV axis, which we defined geometrically by the direction of maximal curvature of the embryo surface.

The degree to which a given PRG stripe deforms over time is dependent upon its position along the AP axis. In general, the closer a stripe is to the posterior pole of the embryo, the more it will deform due to the strong posterior vortices. Therefore, we analyzed the relationship between myosin and PRG orientations on a per-stripe basis (Fig 2j, SI 13), again using Runt as a representative PRG. For Runt stripe three, MRJs are parallel to the DV axis (Figs. 2h,h’,k), in accordance with previous reports [11, 12]. As highlighted by Runt stripe six, in more posterior stripes we observed an increasing myosin/DV axis angle (Figs. 2i,i’,k’). In this region, 22 minutes into GBE, MRJ orientation was less streamlined than at the onset (Fig. 2i’). Quantitatively, we found that the myosin/DV axis angle in Runt stripe three was small throughout GBE, and aligned with MRJs (Fig. 2k). In contrast, in Runt stripe six, which rotated away from the DV axis due to its proximity to the posterior flow field vortex, MRJs reoriented away from the stripe (Fig. 2k’). The global autocorrelation of myosin anisotropy orientation remained consistently high throughout GBE, indicating a nearly stationary pattern (Fig. 2g), akin to instantaneous flow (Fig. 2e).

How to establish an instructive link between a continuously reorienting PRG pattern that moves with the tissue frame of reference, and the nearly stationary direction of both the myosin anisotropy and the flow field is not clear. The dynamic, orientational mismatch is independent of what type of linear or non-linear form of PRG-instructed myosin recruitment is posited. Our observations raise the question: what intra-cellular dynamic rules for myosin recruitment are required to establish the observed global myosin pattern?

### Myosin dynamics are due to tissue-flow driven reorientation and subcellular turnover

To decode the nearly stationary myosin orientation and understand its residual dynamics, we studied the interplay of two dynamic effects: (i) at the subcellular level, the cytoskeleton can dynamically rearrange, due to binding and unbinding of molecular motors to the actin meshwork [24], (ii) at the tissue level, advection will re-orient junctions. We first studied myosin turnover using fluorescence recovery after photobleaching (FRAP). We photobleached individual junctions and measured the signal recovery for about 5 minutes (Fig. 3a). In the first frame after bleaching (1.5 seconds), we measured approximately 50% signal reduction, indicating the presence of a fully mobile myosin subpopulation, possibly cytoplasmic [25]. The recovery curve of *N* = 25 junctions reflected multiple time scales, and had a high standard deviation, possibly due to myosin oscillations on a time scale of ∼60 seconds (Fig. 3a’, Ref. [26]). During the first 30 seconds after photobleaching, fluorescence recovery was rapid, as previously measured [15, 19]. After 30 seconds, the rate of recovery slowed, and pre-bleach levels were reached by 210 seconds, suggesting myosin molecular motors on junctions are dynamically recruited into the cortex and only bind transiently.

**FIG. 3.**
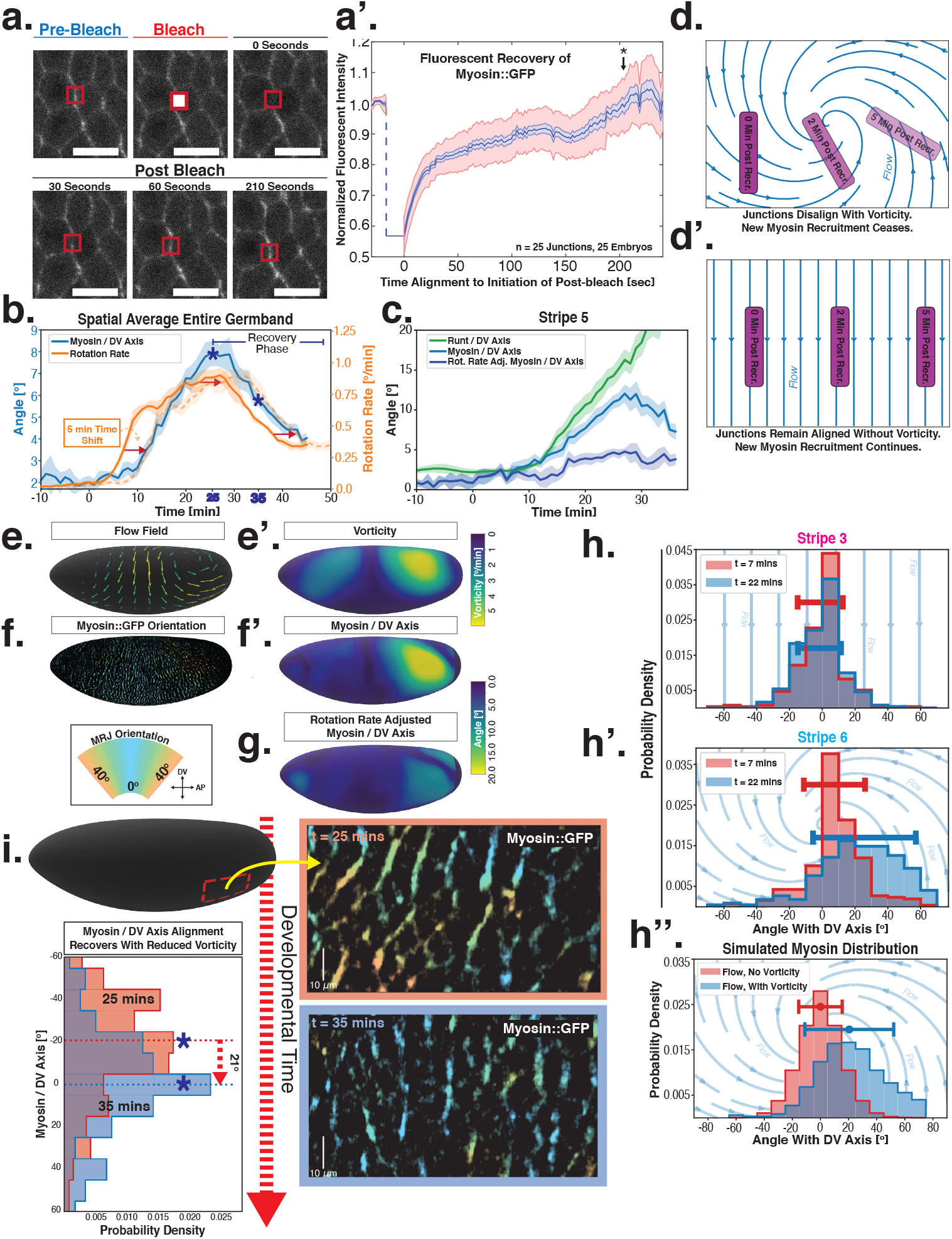
Dynamics of myosin orientation can be quantitatively captured by embryo geometry and vorticity. (a) Time series of a representative fluorescence recovery after photo bleaching (FRAP) experiment of junctional myosin. (a’) FRAP of junctional myosin (*N* =25 embryos) shows multiple timescales and complete recovery of myosin fluorescence, indicating transient binding to the cortex. Rose shaded area indicates standard deviation and blue shaded error the standard error of the mean. (*) indicates the time of full recovery. (b) Spatial average of vorticity and myosin/DV axis angle across germ band versus time in *N* = 5 WT Myosin::GFP embryos. Dashed orange line shows time course of vorticity, shifted by 5 minutes. (c) Spatial average of Runt/DV axis angle, myosin/Runt stripe, myosin/DV axis angle and rotation-rate corrected myosin/DV axis angle over the region corresponding to Runt stripe five versus time in *N* = 5 WT Myosin::GFP and *N* = 5 WT Runt::LlamaTag-GFP embryos. (d-d’) MRJs deflect away from the axis of preferential recruitment in rotational (d) flow but remain aligned in irrotational (d’) flow. (e) Tissue flow field during GBE, temporal average from 15-25 minutes post VF initiation. Computed from ensemble of *N* = 5 WT Myosin:GFP embryos. (e’) Vorticity of tissue flow field, temporal average from 15-25 minutes post VF initiation. Computed from ensemble of *N* = 5 WT Myosin::GFP embryos. (f) MRJs in a representative WT Myosin::GFP embryo 22 minutes post VF initiation. Junction color corresponds to the junction/DV axis angle and junction brightness to the myosin fluorescent intensity. (f’) Smoothed myosin/DV axis angle, temporal average from 15-25 minutes post VF initiation. Computed from ensemble of *N* = 5 WT Myosin::GFP embryos. (g) Rotation rate adjusted myosin/DV axis angle, temporal average from 15-25 minutes post VF initiation. Computed from ensemble of *N* = 5 WT Myosin::GFP embryos. (h-h’) Histogram of myosin angular distribution in the region corresponding to Runt stripe three (H) and stripe six (H’), at two times during GBE. Data corresponds to region shown in Fig. 2h’-i’ (one representative WT Myosin::GFP embryo). (h”) Simulated histograms of Myosin angular distribution the presence or absence of vorticity (rotation rates of 0°*/*min and 3°*/*min, myosin effective lifetime of 5 min.). (i) Junctional myosin in a ventro-posterior region of the embryo at 25 and 35 minutes post VF initiation, showing the recovery of myosin/DV axis alignment. Junctions colored according to their orientation and fluorescent intensity as in (f). Histogram shows the distribution of orientations observed at 25 and 35 minutes.

Next, we characterized the orientation of myosin anisotropy at a tissue scale. The angle between the local myosin anisotropy and the DV axis at every point of the germ band over time, comprises a three-dimensional dataset which we now break down in several ways. We first analyzed the orientation of detected MRJs with respect to the DV axis as a function of time (Fig. 3b). Consistent with the high autocorrelation of myosin orientation (Fig. 2f), we found only a small change in the spatially averaged myosin/DV axis angle. Before the initiation of GBE, the myosin/DV axis angle was 2 degrees. After the onset of GBE flow, measured at 10 minutes post VF initiation, the spatial average increased and reached a maximum of 8 degrees at 25 minutes. Strikingly, after 25 minutes post VF initiation, the average myosin/DV axis angle began to decreases again, as the orientation of myosin anisotropy re-aligned with the DV axis. We refer to this phase of myosin behavior as the “recovery phase”.

We also carried out a regional, stripe-specific analysis to account for the fact that the tissue flow field responsible for advection varies across the germ band (Fig. 3c and SI Fig. 13). Since PRGs advect with cells, we used the angle of individual stripes with respect to the DV axis to characterize local tissue-level reorientation, allowing us to test the degree to which myosin orientation is affected by advection. For Runt stripe five (Fig. 3c), the Runt/DV axis angle was approximately 3 degrees at the beginning of GBE and then increased monotonously starting at 10 minutes, exceeding 20 degrees by 30 minutes post VF formation. The myosin/ DV axis angle measured in the region of stripe five also increased after 10 minutes. This increase was notably less than that of the Runt/DV axis angle from which it clearly diverges by 15 minutes. By 28 minutes, myosin deflection away from the DV axis reached a maximum of 12 degrees, followed by a recovery phase. The same analysis for runt stripe three showed neither of the two quantities accumulated an angle with the DV axis that exceeded 4 degrees (SI Fig. 13).

The two stripes, three and five, are in spatially distinct regions of the tissue flow field. Runt stripe three coincides with the central region of the flow, where instantaneous flow and thus cell trajectories were mainly parallel to the DV axis (Figs. 3d-e). Consequently, we expected the local sense of orientation to be conserved. Runt stripe five was passing through a vortex which we expected to rotate cell junctions (Fig. 3d). To test this idea, we first characterized the rate of local rotation in degrees per minute due to tissue flow (Fig. 3e), by calculating the vorticity (Fig. 3e’) [5]. At the whole embryo level, the spatial pattern of the vorticity had a broad peak of about 5 degrees per minute around the domain of the posterior vortex, and a small peak below 2 degrees per minute on the anterior, vanishing elsewhere. We next characterized the spatial pattern of the orientation of MRJs with respect to the DV axis (Fig. 3f). Across most of the embryo surface, MRJs were parallel with the DV axis, with a clear exception in a domain around the posterior vortex, where individual junctions deflected as far as 20 degrees from the DV axis (Fig. 3f’). This spatial correlation between vorticity and DV axis deflection also extends across time, with a 5-minute delay between the time course of vorticity and the myosin angle defect (Fig. 3b).

### The interplay of subcellular and tissue level dynamics quantitatively captures the myosin pattern

Next, we asked if the vorticity and the myosin/DV axis angle could be causally linked. Myosin stayed bound on junctions for an extended but finite amount of time (Fig. 3a’). We postulated that myosin motors preferentially bind to junctions when they are parallel to the DV axis. Once these junctions are rotated by tissue flow, the myosin orientation is rotated with them, until motors begin to detach (Fig. 3d). The resulting deflection angle is given by the product of rotation rate and myosin lifetime, which we reasoned could quantitatively account for the orientation defect of myosin in regions of high vorticity. We implemented this geometric source hypothesis in a mathematical model that describes the time course of the myosin concentration *m* on a junction with orientation *θ*:

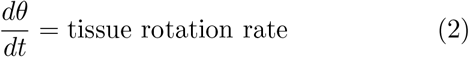

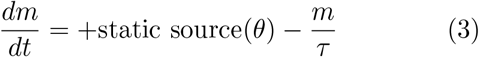

The model assumes that: (i) myosin binds to junctions that are parallel to the DV axis (the “Static Source” is peaked around *θ* = 0), (ii) myosin unbinds with rate 1*/τ*, and (iii) that junctions rotate proportional to local vorticity. Eq. 3 for a single junction can be transformed to describe the entire angular distribution of junctional myosin (see SI Sect. I E) and used to predict the local average deflection angle 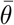 of the myosin anisotropy:

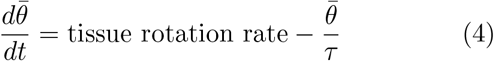

Crucially, 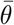 only depends on the direction around which recruitment is maximal and is independent of its magnitude, potential spatial modulation, and functional form (see SI Sect. I E), in agreement with the reasoning presented above. The only free parameter is the effective myosin lifetime *τ*, which captures the duration that myosin motors remain bound on junctions (see SI Sect. I E). Our model makes several quantitative predictions about junction dynamics which we tested (Figs. 3g-i).

First, we computed the angle between the orientation of myosin anisotropy and the DV axis, adjusted for the rate of tissue rotation by solving Eq. 4 (see SI for mathematical definition). We found that junctions aligned well with the DV axis, when the duration *τ* of myosin binding was 5 minutes (Fig. 3g). Assuming this parameter to be constant in time and across the embryo, we found that throughout GBE – even in regions of highest vorticity – the rotation-rate adjusted angle remained close to 0 degrees, while the discrepancy between the Runt and myosin orientations continuously increased (Fig. 3c and SI Fig. 13). Second, we analyzed the angular distribution of MRJs, i.e. the range of orientations of MRJs detected in a small tissue patch, and measured its spread (Figs. 3h-h” and SI Fig. 14). The dynamics of the angular distribution depended on location within the embryo. The standard deviation remained nearly constant in regions of low vorticity, for example around Runt stripe three (Fig. 3h, Fig. 2h-h’). In regions of high vorticity, the standard deviation rapidly changed, giving rise to a much broader distribution (Fig. 3h’, Fig. 2i-i’). This feature is accurately captured in our model: without vorticity there will be no reorientation, and only junctions parallel to the DV axis will recruit myosin (Fig. 3h”). With vorticity, junctions that recruited myosin while aligned with the DV axis will rotate. Since myosin stays bound for a extended but finite lifetime, the distribution widens (Fig. 3h”). Third, we measured the time course of vorticity. It increased with the onset of GBE, reached a plateau by 20 minutes and slowed down by 25 minutes (Fig. 3b). Strikingly, MRJs realigned with the DV axis once vorticity decreased (Figs. 3b,i), as predicted by Eq. 4. The time delay between vorticity and deflection rate is expected from our model: It takes time to deflect junctions in response to vorticity and, correspondingly, for myosin to detach from deflected junctions once vorticity decreases. At 25 minutes, MRJs in a posterior region on the ventrolateral side of the embryo were strongly deflected, with a median angle of about 20 degrees to the DV axis (Fig. 3i). 10 minutes later, the median of the distribution shifted by 21 degrees, aligning with the DV axis. These observations were consistent with the geometric source hypothesis, both qualitatively and quantitatively (Fig. 3c).

Our mathematical model accounts for the dynamics of myosin orientations in terms of the rotation due to flow, and the extended but finite binding lifetime of myosin motors to junctions. The model accurately describes key features of the spatiotemporal dynamics of the mean as well as standard deviation of the distribution of MRJ orientations. The model’s single parameter is the same across the embryo, constant in time, and agrees qualitatively with recovery kinetics as measured on individual junctions experimentally. These results point towards control of myosin orientation by static geometric cues, as opposed to the Lagrangian PRGs.

### Myosin dynamics in genetic and geometric mutants confirms static orientation of myosin recruitment

Harnessing the genetic toolkit available in the *Drosophila* model system, we can independently modulate parameters of our mathematical model – vorticity, myosin kinetics, and geometry (Figs. 4a-a”). Twist is expressed in the VF and *twist*^*ey*53^ mutants have a defect in VF formation, accompanied by reduced kinetics across the entire embryo [23, 26–28]. We found the average speed of GBE in *twist*^*ey*53^ mutants was reduced by a factor of two compared to WT (Fig. 4b). This was accompanied by a corresponding reduction of the vorticity (Figs. 4a, c). The myosin/ DV axis angle was likewise smaller (Figs. 4a, c). Fitting the model to this data revealed a similar myosin binding lifetime as in WT, and the rotation rate adjusted angle of MRJs closely aligned with the DV axis.

**FIG. 4.**
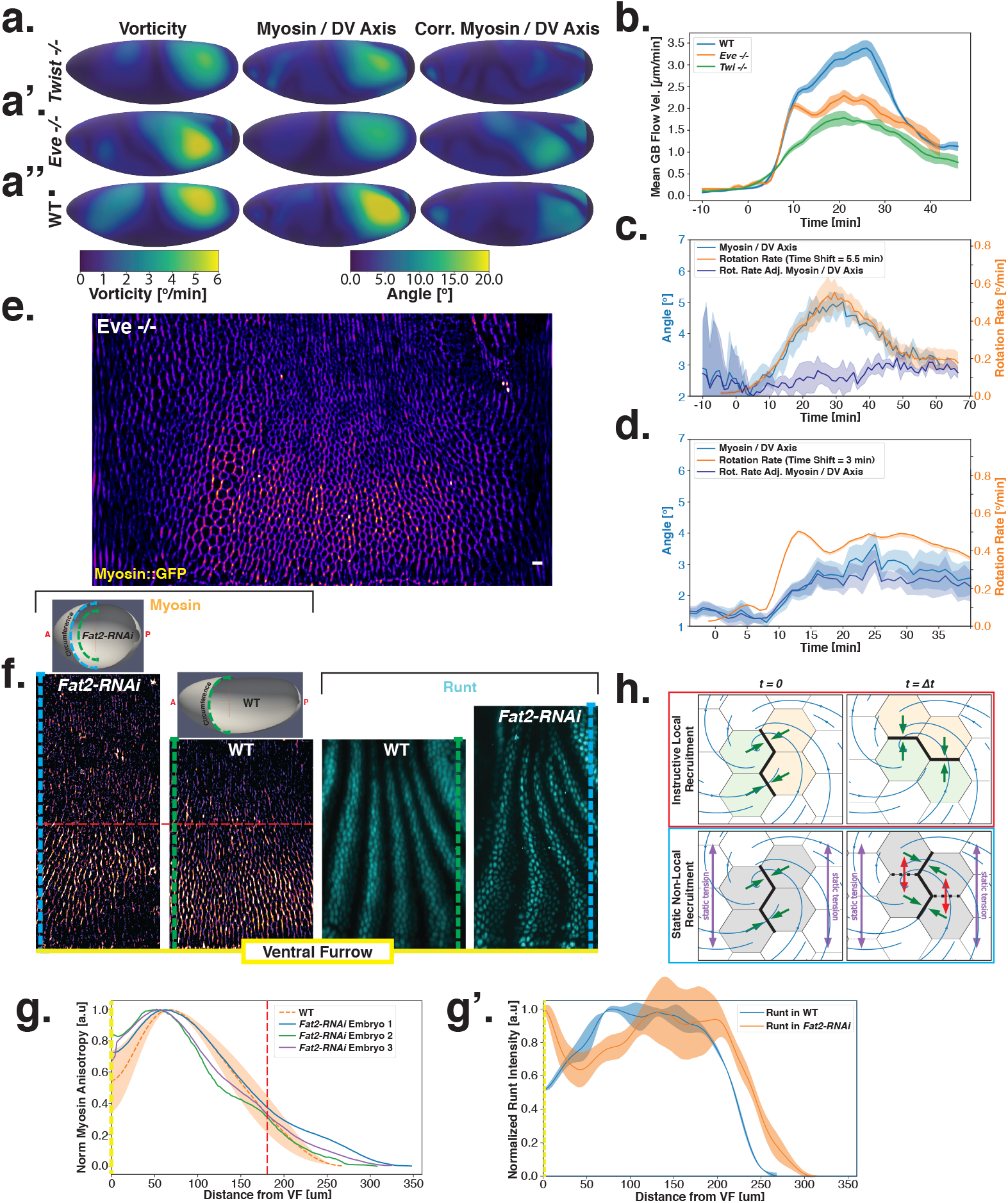
Dynamics of myosin orientation in mutants affecting vorticity or embryo geometry can be quantitatively described. (A-A”) Vorticity, myosin/DV axis angle, and rotation-rate adjusted myosin/DV axis angle prediction in *N* = 5 WT, *N* = 4 *twist*^*ey*53^, and *N* = 3 *eve*^*R*13^ embryos. Heatmaps show a temporal average from 15-25 minutes post VF initiation. The correlation of vorticity and myosin/DV axis angle persists in all mutants, and the rotation-rate corrected myosin/DV axis angle remains low. (B) Spatial average of tissue flow velocity in the germ band over time in *N* = 5 WT, *N* = 4 *twist*^*ey*53^, and *N* = 3 *eve*^*R*13^ embryos. *twist*^*ey*53^ and *eve*^*R*13^ mutants show markedly lower flow velocity than WT, but differ in their kinetics. (C) Spatial average of vorticity, myosin/DV axis angle, and rotation-rate adjusted myosin/DV axis angle over time in *N* = 4 *twist*^*ey*53^, Myosin::GFP embryos. (D) Spatial average of vorticity, myosin/DV axis angle, and rotation-rate adjusted myosin/DV axis angle over time in *N* = 3 *eve*^*R*13^, Myosin:mCherry embryos. (E) Junctional myosin in *eve*^*R*13^ mutants remains anisotropic and aligned with the DV axis, although the degree of anisotropy is reduced. One lateral half of a representative *eve*^*R*13^, Myosin::mCherry embryo, 18 min. post VF initiation. (F) Top: 3D-shape of WT embryos and embryos from *Fat2-RNAi* mothers, extracted by tissue cartography pipeline. Compared to WT, *Fat2-RNAi* embryos are spherical and have a greatly increased circumference (marked in green resp. blue). Bottom: Patterns of junctional myosin and Runt in the germ band of WT and *Fat2-RNAi* embryos at equivalent phases in GBE (10 min post VF initiation in WT). Only one lateral half is shown. Junctional myosin is visible up to the same distance from the VF in both WT and *Fat2-RNAi* embryos. (G-G’) Quantification of the decay of junctional myosin (G) and Runt (G’) away from the ventral furrow in WT and *Fat2-RNAi*. Myosin data from *N* = 5 WT and *N* = 3 *Fat2-RNAi* embryos. Runt data from both lateral halves of WT and *Fat2-RNAi* embryo shown in (F). (H) Myosin recruitment by a Lagrangian source vs by a static source leads to qualitatively and quantitatively different behavior.

Analysis of fixed samples has demonstrated that during GBE, myosin polarization in PRG and TLR mutants is reduced [12]. GBE tissue flow in these mutants is impaired as well, particularly in later phases [10]. Our dynamic data indicates that myosin and PRGs/TLRs are regulated in different frames of reference. This raises the question of how patterning gene expression can be quantitatively linked to myosin anisotropy and – by extension – tissue flow. To study this, we performed live imaging of myosin in *eve*^*R*13^ mutants (Figs. 4a’,b,d,e). We found that the initial kinetics of GBE closely match that of WT embryos. However, at 10 minutes post VF formation the kinetics of *eve*^*R*13^ mutants change abruptly (Fig. 4b). Additionally, vorticity was reduced compared to WT (Fig. 4d). We detected anisotropic myosin in the germband during GBE (Fig. 4e), although anisotropy was significantly reduced in comparison to WT (SI Fig. 16). We found that MRJs were mainly aligned with the DV axis, except in the region of the posterior vortex (Fig. 4a’). The best fit of the static source model to this data suggests that myosin binding lifetime is significantly reduced in *eve*^*R*13^ mutants (to *τ* = 2 − 3min). A joint reduction of vorticity, and myosin lifetime accounts for the near constant time course of the MRJs angle defect with the DV axis (Fig. 4d).

Finally, by knocking down the atypical cadherin Fat2 (*Fat2-RNAi*) in somatic ovarial cells in female flies, we created nearly spherical embryos [29] with a shorter, but variable length, and up to 30% extended DV circumference (Fig. 4f, top). This provided an opportunity to test if PRG stripes would change in the same way as MRJs around the ectopically extended DV circumference. The AP patterning system remains intact in *Fat2-RNAi* embryos [29]: PRGs were expressed in seven stripes along the AP axis, with a decrease of expression at the dorsal pole (Fig. 4f, right). Similar to WT, myosin recruitment to junctions was strongest in ventral regions, and dropped markedly on the lateral side. In both WT and *Fat2-RNAi* embryos, MRJs were detected up to ∼175 *μ*m away from the VF in the lateral ectoderm (Fig. 4 g-g’). Strikingly, since the absolute length of the DV circumference is larger in *Fat2-RNAi* embryos than it is in WT embryos, and the PRG stripes extended normally to the dorsal pole, there is a substantial region on the lateral surface of *Fat2-RNAi* embryos where PRG stripes are clearly visible but no myosin anisotropy could be detected (Fig. 4f). The magnitude of tissue flow in *Fat2-RNAi* was reduced (SI Fig. 17). Moreover, MRJs orientation changed little, due to low spatial overlap between the regions of high vorticity and myosin recruitment (SI Fig. 18). These observations suggest that presence of striped PRG expression is not sufficient to set up myosin anisotropy.

Taken together, these results suggest that instead of instructing anisotropic myosin recruitment, PRGs influence the myosin anisotropy by regulating retention of myosin to junctions.

## DISCUSSION

Here we presented a quantitative study dissecting the dynamic rules governing myosin anisotropy during *Drosophila* GBE. We found that the orientation of MRJs closely tracks the DV axis, a static geometric landmark. By contrast, the localization of patterning genes (PRGs and TLRs) implicated in GBE, are Lagrangian. They deform due to advection with the flowing tissue, and deflect away from the DV axis over time (Fig. 4h). We define a mathematical model which accounts for the dynamics of myosin orientation as a product of tissue flow vorticity and the extended-but-finite time that myosin motors remain bound to junctions.

These results suggest that the known upstream regulatory factors of GBE – PRGs and TLRs – are Lagrangian and follow a different reference frame than myosin. This observation makes it difficult to posit a direct, instructive link between anisotropic myosin recruitment and local differences in PRG levels between adjacent cells. We show that such a connection would be complex and non-linear. Results from *Fat2-RNAi* embryos further indicate that the presence of PRG stripes is not sufficient for anisotropic myosin recruitment. Our model presents a simpler alternative with only a single parameter and suggests a clearly interpretable biophysical role for PRGs (likely via the TLRs): modulating the sensitivity of myosin recruitment to the static source and regulating myosin maintenance on junctions.

Our dynamic data from WT as well as multiple mutant genotypes are consistent with preferential myosin recruitment along the DV axis. However, the mechanism underlying myosin recruitment remains unclear. Myosin dynamics can be organized not only by instructive genetic signals but also by mechanical inputs [15, 26, 30]. Crucially, mechanical cues such as epithelial tension are not necessarily advected by tissue flow.

Strain-responsive myosin recruitment, triggered by DV strain due to the invagination of the VF, establishes early myosin anisotropy, acting as starting signal for GBE [26]. Yet VF formation is transient, raising the question of how anisotropic recruitment is maintained during later stages of GBE. One stationary signal with the required anisotropy is mechanical feedback triggered by epithelial stress [19, 31, 32]. The static stress anisotropy might originate from turgor pressure within the embryo [33], which the surface stress needs to balance [34]. Due to the cylinder-like geometry of the embryo, this results in a static, anisotropic surface stress (“hoop stress”) [35]. Cortical tension due to turgor pressure is known to play a crucial role in mouse blastocyst development [29]. Tools for faithful measurement and manipulation of hoop stress will be needed to further evaluate this hypothesis.

The geometric control of myosin orientation described here has close parallels to other model processes of axis elongation, such as the *Drosophila* wing disc [36] and the *Xenopus* larval epithelium [37]. In both cases, planar cell polarity proteins orient according to mechanical inputs propagated over tissue-length scales. Our results suggest that underlying biological complexity notwithstanding, the dynamics of morphogenesis can be quantitatively described by simple models with few and clearly interpretable parameters.

## ACKNOWLEDGMENTS

The authors thank Eric Wieschaus, Boris Shraiman, Fridtjof Brauns, and members of the Streichan lab for valuable discussions and suggestions. We additionally wish to thank Eric Wieschaus for providing several fly lines, Sophie Streichan for aid in handling stocks, crosses, and reagents, and Cécile Regis for assistance with 3d visualizations. This research was supported by NIH grant No. R35 GM138203 and partially supported by NSF grant PHY–1707973. MFL acknowledges support by NIH F31 fellowship No. HD093377. NPM acknowledges support from the Helen Hay Whitney Foundation.

## MATERIALS AND METHODS

### Lightsheet Microscopy

#### Microscopy

Lightsheet data sets were taken on a custom Multi View Selective Plane Illumination Microscope (MuVi SPIM) [17] with scatter reduction through confocal imaging [38]. This microscope is capable of fluorescent imaging of the entire *D. Melanogaster* embryo at subcellular resolution and was previously described in detail in Ref. [26]. Electronics were controlled using MicroManager [39].

#### Image acquisition

Prior to imaging, embryos were dechorionated and mounted in low-melting point agarose gel [17]. Samples are imaged simultaneously by two objectives at opposite sides of the embryo, with lighsheet *z*-sections spaced by 1.5 *μ*m. By rotating the embryo by 45°, 90° and 135°, and repeating the *z*-imaging, we create a total of 8 views per time point which are registered and fused to create a volumetric dataset in the next step. All lightsheet movies in this work are taken at a time resolution of 1 minute.

#### Data fusion and surface extraction

Images recorded by the lightsheet microscope were registered based on the position of fiduciary beads embedded in the agarose (Fluoresbrite multifluorescent 0.5-*μ*m beads 24054, Polysciences Inc., as described in Ref. [26]) using the Multiview reconstruction plugin [40] in Fiji [41]. We used all-to-all registration, mapping all perspectives at all time points to a common reference frame using an affine transformation. Images were then deconvolved and fused using the algorithm introduced in Ref. [40], yielding images with an isotropic resolution of 0.2619*μ*m.

The embryo surface is detected within the resulting volumetric data using an Ilastik detector [42], to which a surface was fitted using the ImSAnE software [18] which was used for tissue cartography as described in Ref. [18]. To improve accuracy, we applied two iterations of the Ilastik+ImSAnE workflow. The resulting “onion” layers normal to the embryo surface, spaced 1.5 *μ*m, were used to generate maximum-intensity projections.

### Confocal Imaging and FRAP

Imaging for the FRAP experiments shown in Fig. 3 and the *eveeve*^*R*13^ data shown in SI Fig. 15 was done using a Leica SP5 confocal microscope and 63x/1.4 NA oil immersion objective at a frame rate of 1 frame per 1.78 seconds. Junctions were tracked manually using Fiji Freeware Software [41]. We bleached regions of size 5*μ*m × 5*μ*m for approximately 8 seconds, using 50mW laser power.

### Fly stocks and genetics

A full stock list is presented in Table S1. The fluorescent fusion proteins used in this study include: Myosin::GFP (II or III, sqh::GFP, [43]), Myosin::mCherry (II, sqh::mCherry, [27]), Runt::LlamaTag-GFP (, [44], gift from H. Garcia), Eve::YFP (III, [45]), Gap43::mCherry (III, Membrane::mCherry, [46]).

Recombinant chromosomes containing the chromosomal deficiency Df(2L)dpp[s7-dp35] 21F1–3;22F1–2 (*halo*) and either *eve*^*R*13^, or *twist*^*ey*53^ were balanced with CyO. The chromosome containing *eve*^*R*13^ was recombined with Myosin::mCherry (II). *Halo, twist*^*ey*53^ embryos were balanced with a version of CyO that also contains Myosin::GFP. Homozygous *eve*^*R*13^ and *twist*^*ey*53^ embryos were identified based on visualization of the *halo* phenotype while hemizygous control embryos lacked *halo*.

The following stocks were used to generate reduced aspect ratio (*Fat2-RNAi*) embryos: w; Traffic jam-Gal4; Myosin::GFP; Gap43::mCherry, w; Myosin::GFP; UAS-*Fat2-RNAi* [47].

### Immunohistochemistry and Antibody Production

For heat fixation, embryos were dechorionated with 50% bleach and then fixed using heat and methanol as described previously [48]. Primary antibodies for immunohistochemistry were Runt (Guinea Pig, 1:500, Wieschaus Lab), Even-Skipped (Rabbit, 1:500, gift from M.Biggin), Fushi-Tarazu (Rabbit, 1:1000, Gift from M.Biggin), Paired (Mouse, 1:100, Gift from N.Patel), Sloppy-Paired (Rabbit, 1:500, Gift from M.Biggin), Hairy (Rat, 1:100, Wieschaus Lab), Tartan (Rabbit, 1:100, This study, GenScript, based on Full-length peptide). Donkey and Goat secondary antibodies conjugated to AlexaFluor488, 561, and 647 were used (1:500, Thermo Fisher Scientific). Embryos were mounted in 1.5% low gelling temperature agarose (Millipore Sigma-Aldrich) for light-sheet imaging, and mounted in 50%PBST 50%Aqua-Poly/Mount (Polysciences) for confocal imaging.

### Image processing and analysis software

Image processing, described in detail in the SI, used a combination of custom python scripts using the scientific python [49] and scikit-image [50] packages and custom MATLAB scripts. Tissue cartography was performed using the ImSAnE software [18]. Surface detection, cell tracking, and segmentation of Runt stripes was performed using Ilastik [42].

### Data availability

All data and analysis code not contained in this article is available from the authors upon reasonable request

## SUPPLEMENTARY INFORMATION

### A. Quantitative image analysis

#### 1. Tissue cartography and image analysis on curved surfaces

To analyse the *in toto* 3d data obtained by light-sheet microscopy [17], we use tissue cartography as described in Ref. [18]. Briefly, the embryo surface is detected using a machine-learning pixel classification workflow and a smooth surface is fit to the resulting point cloud. This surface is refined in a second step, repeating classification and fitting. This surface can be evolved along its normal to create so-called onion layers. We created pullbacks showing the pixel intensity on these onion layers and obtained the final image by a maximum projection along across layers.

The result shows the embryo in a cylindrical chart with anterior left and posterior right. The ventral midline corresponds to the cut used to unroll the cylinder. Velocity fields, gradients of fluorescent intensity, and orientations of cell edges can then be computed within this chart. All angles are calculated with respect to the induced metric, correcting for distortions induced by the cylindrical projection near the poles.

For *Fat2-RNAi* embryos, which have a particularly round shape, we also used an second cylindrical projection, analogous to the Mercator projection of the earth, which is angle-preserving at all points. This allows our edge-detection algorithm (see below) to work faithfully and avoids spurious anisotropy detection near the poles. The detected edges were then mapped back to the original cylindrical chart.

#### 2. Time alignment and dynamic atlas

In our work, we combine and compare data from different embryos carrying with complementary makers. Note that because we employ *in toto* imaging and tissue cartography, spatial alignment is trivial. Temporal alignment is facilitated by the extreme reproducibility of the movements of gastrulation across embryos. Embryos were time-aligned using two methods.

##### a. Time alignment of fixed embryos

Fixed embryos were time-aligned using a landmark based approach. In addition to its primary staining, each fixed embryo way stained for Runt, and the anterior boundary of the seventh Runt stripe was computationally extracted. The same was done for *N* = 5 live recordings of embryos with fluorescent tagged Runt. Live movies were time aligned to one another using all-to-all optimization of the similarity of the extracted stripe contour across movies. The stained samples were aligned to the resulting master time line. The details of this “dynamic atlas” method will be described in a forthcoming paper (Mitchell et al. 2022).

##### b. Time alignment of live data

To time-align different live films not tagged for Runt, we compared their PIV-calculated flow fields as in Ref. [23]. PIV fields were calculated using the phase-correlation method as in Ref. [23]. For each genotype, we chose one reference movie to which the remaining movies are aligned with a constant time shift *t*_off_, obtained by minimizing the average difference of the velocity fields:

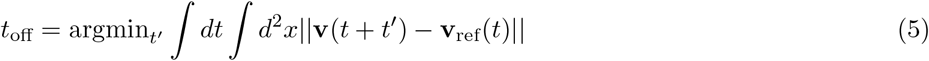

Here, **v** is the velocity field of the embryo to be aligned, **v**_ref_ that of the reference movie, and the integral represent averages over time and the embryo surface. Fig. 5 shows the aligned the spatial average velocities of *N* = 5 WT Myosin::GFP embryos over time. Fig. 5 highlights that time-alignment is possible within ±1min due to the highly reproducible time-courses.

**FIG. 5.**
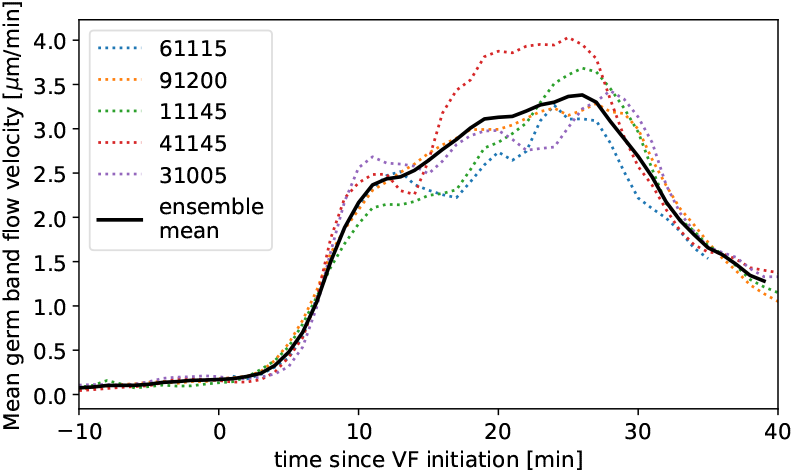
Time-alignment using PIV curves. Average flow velocity in germ band during GBE in *N* = 5 WT Myosin::GFP, time-aligned by matching PIV fields.

#### 3. Ensemble averages and meso-scale analysis

Using the time-alignment obtained, we can compute ensemble-averages across embryos, such as the average velocity field or the average myosin anisotropy. The ability to faithfully compute ensemble averages and thus distill the behavior of the stereotypical embryo and make statistically significant statements is a key advantage of our workflow which combine *in toto* tissue cartography, time-alignment, and mesoscale analysis.

In meso-scale analysis, we smooth cell-level quantities over the scale of ∼3-5 cells to obtain everywhere-defined tissue-level fields [23]. This process drastically reduces noise, focuses on the tissue-level dynamics relevant for large-scale morphogenesis, allows easy comparison across embryos, and defines suitable inputs for the type of quantitative, predictive model we study in this paper. Examples of this approach are the local myosin anisotropy orientation, the smoothed PRG gradients, and the PIV-computed tissue flow field.

We use the ensemble velocity field of *N* = 5 WT Myosin::GFP embryos for the calculation of the vorticity in Fig. 3 and for the transport of Runt stripes in Fig. 2. To time-align Runt and myosin data, we align the Runt PIV fields to the myosin ensemble field according to the onset of GBE flow.

#### 4. Quantitative analysis of junctional myosin

##### a. Cytosolic normalization of junctional myosin

We measure the concentration of fluorescent myosin motors in reference to the concentration of motors in the cytoplasm, an approach validated in Ref. [26]. The cytoplasmic intensity *I*_*c*_ is calculated by applying a top-hat transform with a disk-shaped structuring element with a 1-cell-diameter radius to the image data, intensity *I*. The cytosolically normalized signal is

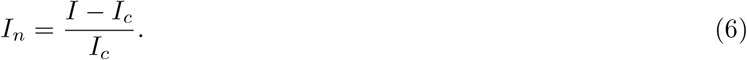

This measure is in principle independent of the concentration of fluorescently tagged molecules and allows to compare data from fly lines with different fluorescent tags. The cytoplasm acts as a pool from which motors can be recruited to the actomyosin cortex. If no motors have been recruited, the concentration near the membrane and in the cytosol will be equal, and the normalized signal, which measures excess junctional accumulation, vanishes.

Fig. 6 shows the average junctional myosin level on junctions in WT and *twist*^*ey*53^ mutants over time. As previously reported [26], junctional myosin levels in *twist*^*ey*53^ embryos are much lower than in WT due to slower recruitment during early GBE. This is a consequence of the lack of VF invagination in which generates forces that drive mechanosensitive myosin recruitment in WT.

**FIG. 6.**
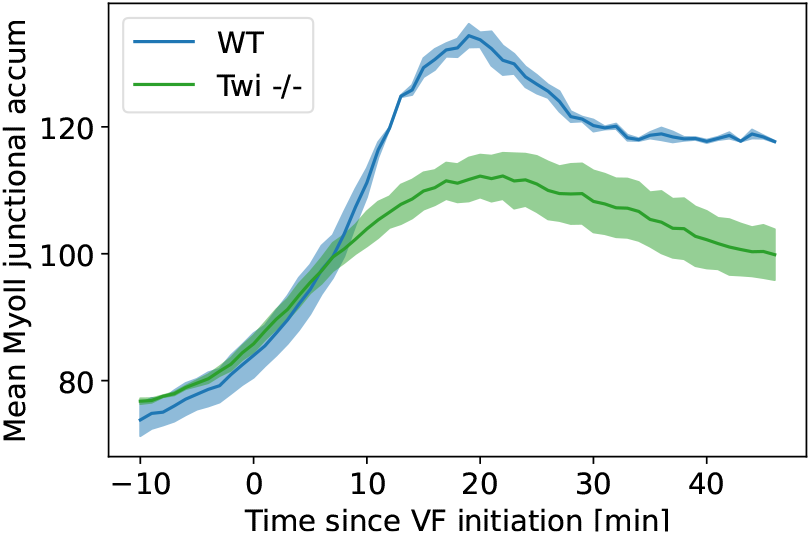
Average junctional myosin levels in WT and *twist*^*ey*53^ embryos. Average junctional myosin levels, computed using the cytosolic normalization filter, in germ band during GBE in *N* = 5 WT Myosin::GFP and *N* = 4 *twist*^*ey*53^ Myosin::GFP embryos. Included in the spatial average are image regions with a junctional accumulation level of ≥.05 which we classify as “junction”.

##### b. Segmentation-free edge detection by Radon transform and computation of local myosin orientation

Cell segmentation at the whole-embryo scale is time consuming. Additionally, for anisotropically distributed markers such as myosin, it is often difficult detect cell edges with low marker levels and obtain correct cell outlines. We therefore used a segmentation-free technique previously presented and validated in Ref. [23]. Briefly, the image is scanned with a local edge detection filter which analyzes circular patches of ∼1 cell diameter at a time. The filter applies the radon transform which computes the normalized line integral of the signal as a function of line orientation 0 ≤ *θ* ≤ 180° and offset of the line from the image center. In this way, edges are mapped to peaks in radon plane whose angle and offset correspond to edge orientation and real-space position and whose heights to the average image intensity along the edge. Such peaks can be detected robustly with different methods. Here, we consider only the global maximum in the radon plane since there is typically only a single edge in the filter window, additionally subject to a criterion filtering out insignificant peaks (peak elevation greater 1.5× the average). Alternative methods, e.g. the h-maxima transform, lead to equivalent results.

This method results in a list of edges containing their positions, orientations, and intensities. From this, we can compute the local average myosin anisotropy orientation, as well as the standard deviation of the orientation distribution. Since angles are defined only modulo 180° (e.g. an edge of angle 5° and 175° are in fact very close), directly averaging the angles can lead to distorted results. We therefore defined a nematic tensor *m*_*e*_ for each edge [51]:

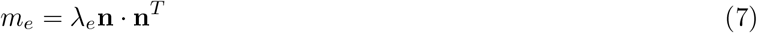

where *λ*_*e*_ is the average fluorescent intensity on the edge and **n** is a unit vector parallel to the edge. These tensors can be locally averaged over a scale of *σ* ≈ 5 cells to produce a tensor *m* representing the local myosin anisotropy. The top eigenvector of *m* defines the local myosin orientation used in Fig. 2-4, and the top eigenvalue *λ* is the average magnitude of junctional myosin on edges aligned with the local orientation.

##### c. Definition of correlation coefficient for nematic fields

For nematic fields, such as the myosin tensor Eq. 7 and the Runt line field, the orientation angle is defined only up to 180°. We therefore use the following definition for the correlation between two nematic directors **n, m**, in particular in the correlation matrices Fig. 2B:

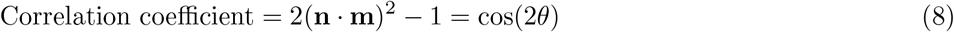

where *θ* is the angle between the directors.

#### 5. Definition of Runt stripe angle

The local orientation of the Runt stripes was calculated as follows:

- Smooth the Runt fluorescence field *ϕ*(*x, y*) with a Gaussian kernel 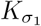 of width *σ*_1_ ≈1 cell diameter
- Compute the smoothed gradient 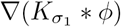, and normalize its magnitude
- Construct the local nematic tensor 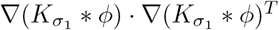
- Smooth the nematic tensor further by *σ*_2_ ≈ *σ*_1_, so that the director is defined over the entire embryo scale (interpolating between stripes)
- The local Runt orientation is defined by the dominant eigenvector of the resulting tensor field.

### B. Additional Data on PRG stripe, TLR stripe, and myosin orientation

In this section, we collect some additional data on the behavior of the genetic pattern of PRGs and TLRs as well as on the myosin orientation. Fig. 7 shows a quantification between the angle of the PRG Runt and the TLR Tartan, complementing Fig. 2A in the main text. Fig. 8 shows that the PRG Eve is advected by tissue flow in the same way as Runt (see main text Fig. 2A). Fig. 9 shows single-cell tracking data confirming the Lagrangian behavior of Runt.

**FIG. 7.**
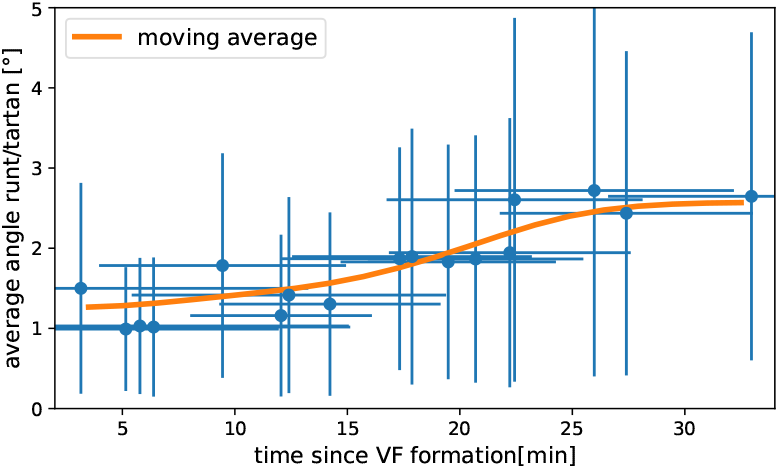
Measured angle between Runt and Tartan stripes. Runt angle from *N* = 5 Runt::LLamaTag-GFP embryos, Tartan data from *N* = 17 stained embryos.

**FIG. 8.**
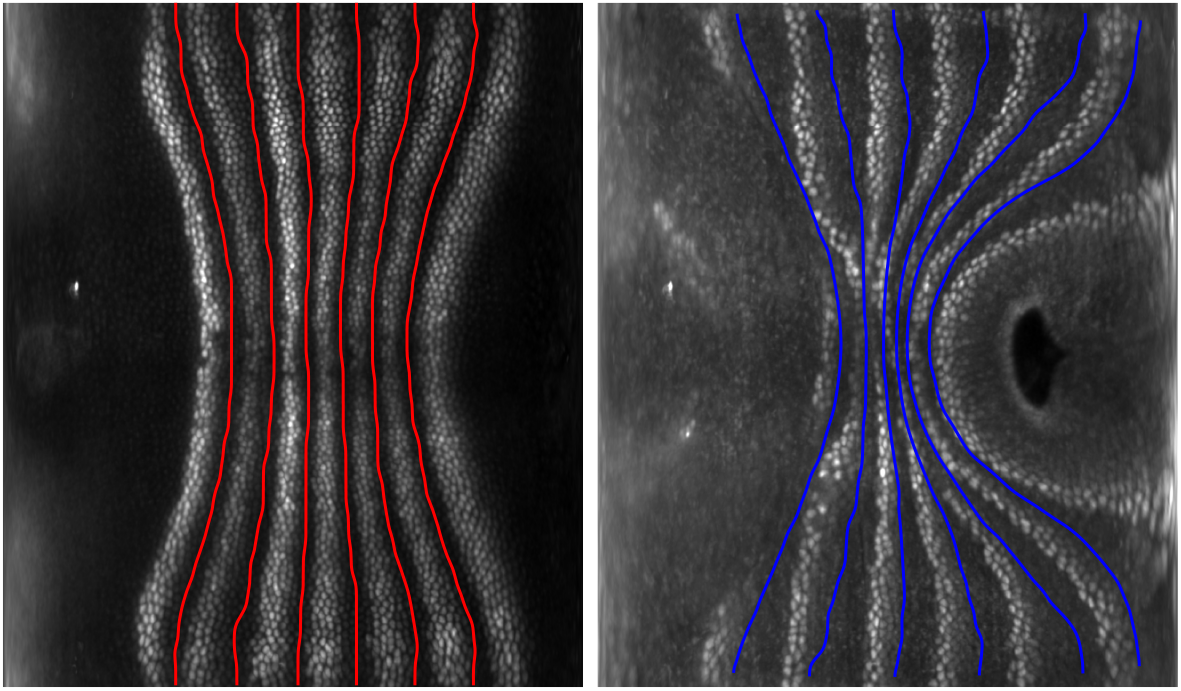
Eve stripes are advected by tissue flow. Eve stripes 5 minutes before VF inititaion (left), and Eve stripes 20 minutes post VF initiation, with predicted inter-stripe locations based on advection (right).

**FIG. 9.**
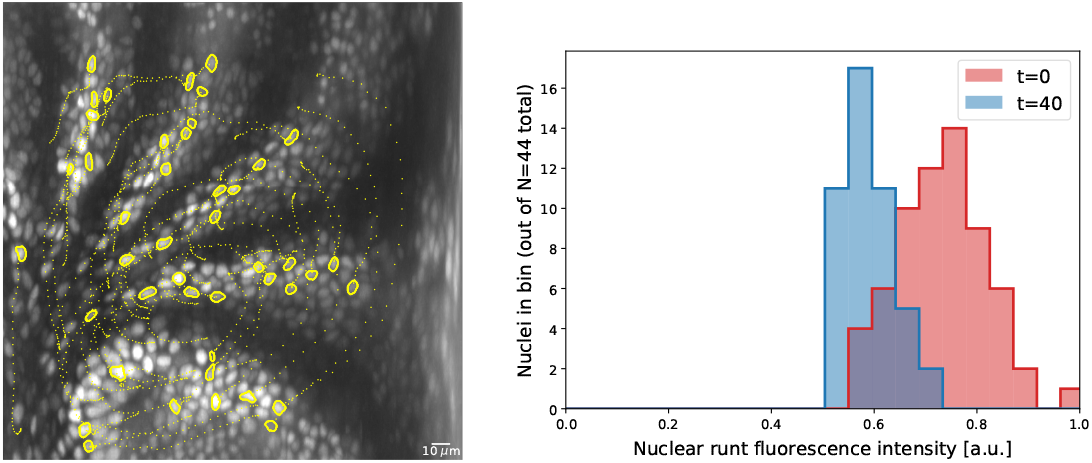
Nuclei initially expressing Runt expression after 20 minutes. Cell tracks obtained by semi-automatic cell tracking in Ilastik.

Figs. 10, 11, and 12 show additional data on the local angular distribution of myosin and Runt (see also SI Fig. 14). Fig. 10 complements Fig. 2F by showing histograms of both the local myosin and the Runt orientation in the ventro-lateral of the germ band. In comparison to Fig. 2F, where for simplicity all myosin junctions in the region under investigation were combined in a histogram, here, we proceeded in a two step fashion. The region was scanned with a 30*μ*m × 30*μ*m window, for each window the mean orientation was calculated and subtracted from the angles of the junctions in the window, and the resulting distributions were joined. This process permits separating local spread from larger scale spatial gradients. Fig. 10 also shows the corresponding distributions for the Runt stripe orientations in the same region. Not only is the spread much smaller, the distribution also changes much less. The total spread of the Runt stripe distribution depends on the smoothing parameters used to compute the Runt gradient, but the dynamics of the distribution does not. Fig. 11 shows the spatial pattern of the local Runt angular distribution across the germ band. Note the strong dissimilarity with the corresponding figure for myosin, Fig. 14. Finally, Fig. 12 carries out a embryo-scale comparison of the local Runt and myosin distributions, as illustrated in Fig. 10, using the entire ensemble of 10 embryos available to us. We compute the Kolmogorov–Smirnov, a measure of distance between two probability distributions (the maximal area between the two probability densities), of the local angular distributions across the germ band, finding high values strongly indicative of disagreement between the two. Each window typically contains more than 100 junctions.

**FIG. 10.**
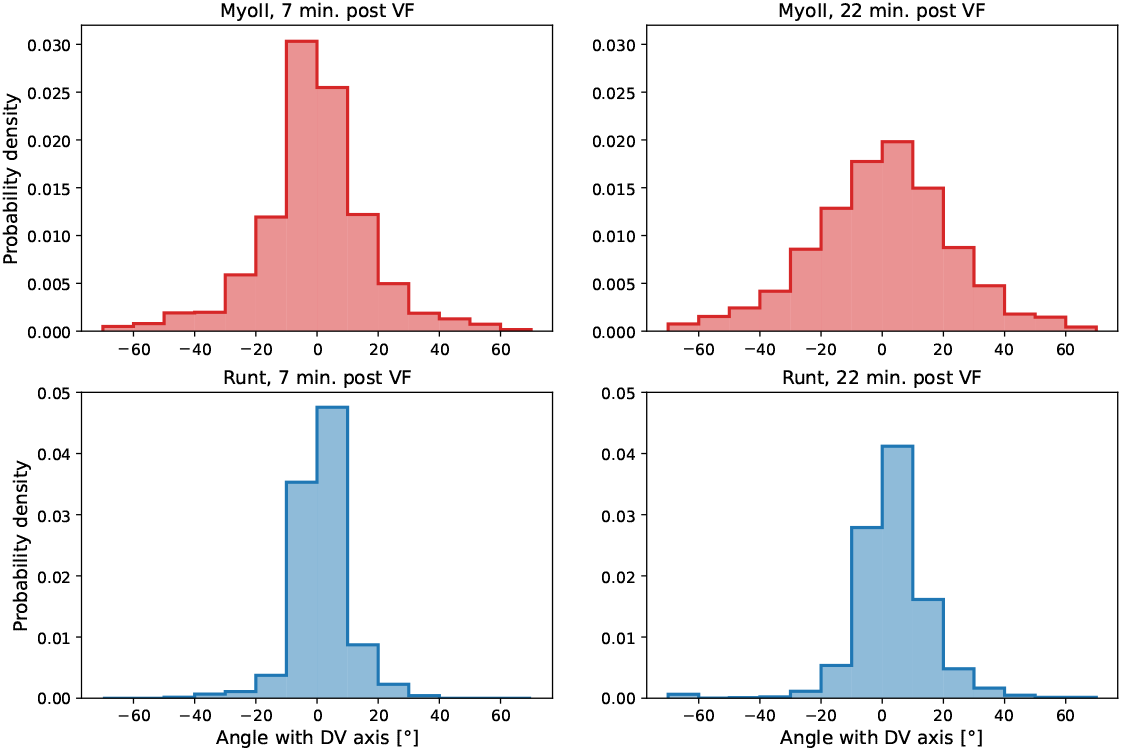
Local distribution of Runt and myosin orientations in the ventro-lateral region of two representative embryos 22 minutes post VF initiation. Runt angle and myosin data taken from to the regions shown in Fig. 2E’. Window size used for querying the local distribution is 30*μ*m × 30*μ*m.

**FIG. 11.**
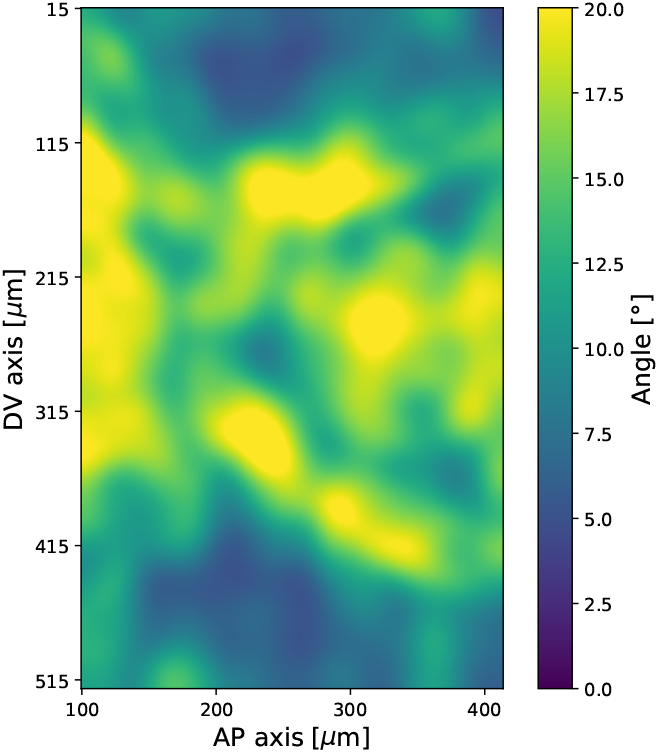
Spatial pattern of local standard deviation of Runt orientation, 20 minutes post VF initiation. Data from *N* = 5 Runt::LlamaTag-GFP embryos. Window size used for querying the local distribution is 30*μ*m × 30*μ*m.

**FIG. 12.**
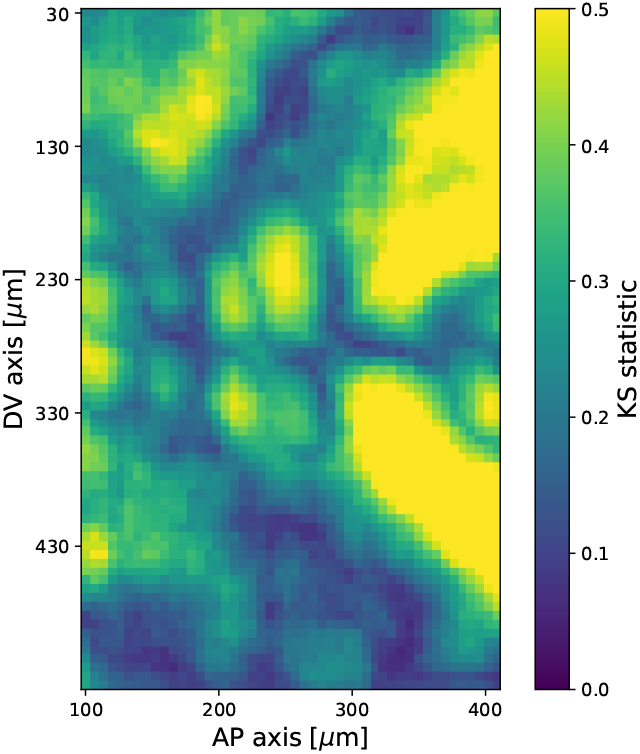
Kolmogorov–Smirnov test statistic comparing local angular distributions of Runt and myosin 20 minutes post VF initiation. Observed values of the test statistic strongly indicate that the two distributions are dissimilar. Runt angle from *N* = 5 Runt::LLamaTag-GFP embryos, myosin data from *N* = 5 Myosin::GFP embryos. Each local window contains ∼100 junctions.

Fig. 13 completes the argument of main text Fig. 3 by showing the time course of PRG and myosin orientation as well as our prediction for the myosin orientation in the regions of all 7 Runt stripes.

**FIG. 13.**
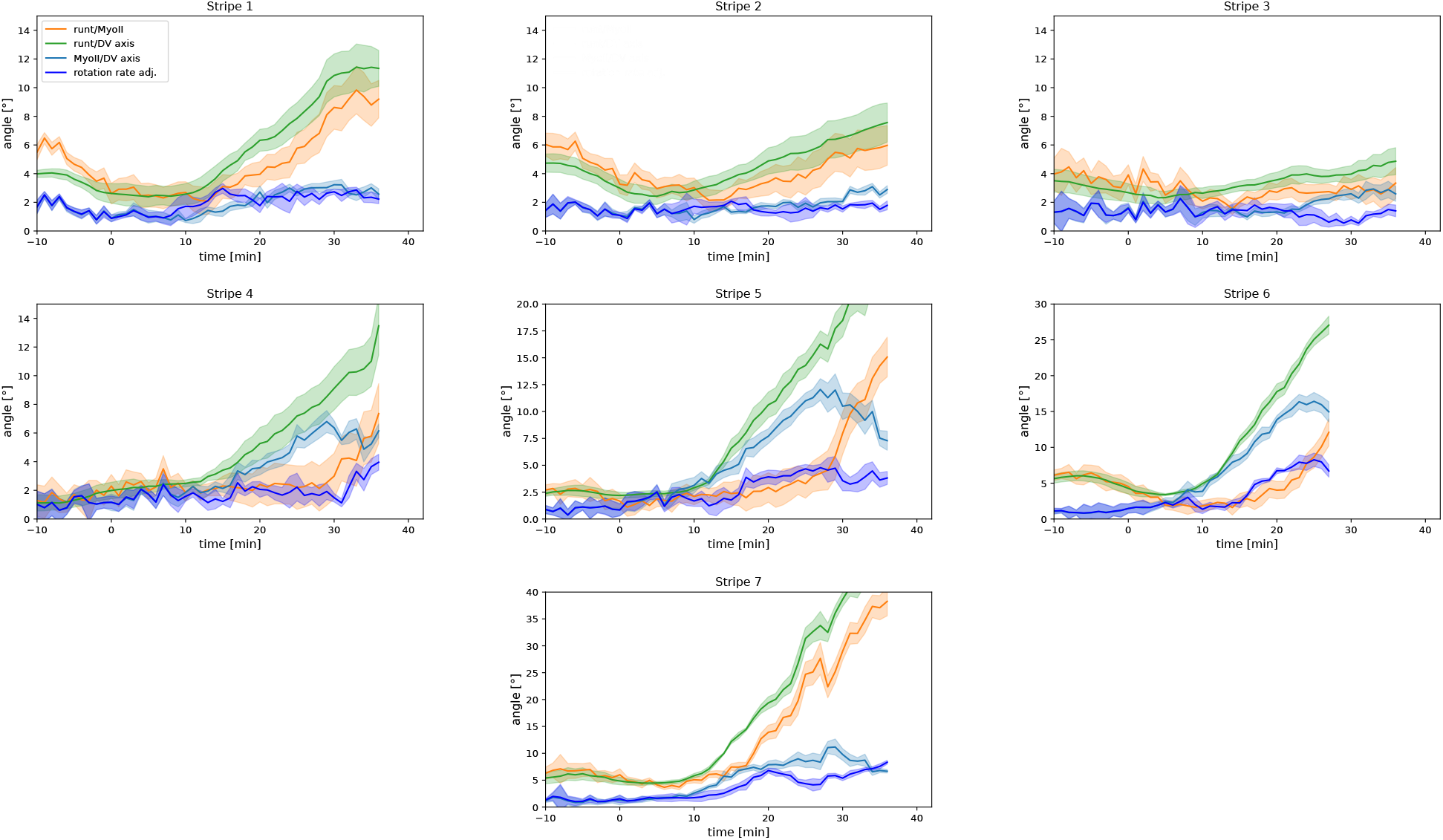
Per-Runt stripe averages of myosin and Runt orientations. Angle between myosin anisotropy orientation and Runt stripe, angle between Runt stripe and DV axis, angle between myosin anisotropy and DV axis and rotation rate adjusted myosin/DV axis angle, averaged over the regions corresponding to Runt stripes 1-7. Data from *N* = 5 WT Myosin::GFP and *N* = 5 Runt::LlamaTag-GFP embryos. Data ends early in stripe 6, since the region becomes difficult to separate from stripe 7.

### C. Normalization of FRAP data

To obtain the FRAP curves show in Fig. 3 from the raw recorded data, we used a two-step normalization procedure. The average raw intensity in the region of interest (ROI) containing the bleached junction will be denoted *I*_*t*_. *t* denotes time, with *t* = 0 being the first frame after bleaching. The bleached region is monitored for a total time *T*. In addition to the ROI containing the bleached junction, we also monitor three additional 5*μ*m × 5*μ*m control regions not containing any junctions. Their average defines the background intensity *I*_bg,*t*_.

In a first step, we normalize the ROI intensity by the background, which also corrects for photo-bleaching, defining *Ĩ*_*t*_ = *I*_*t*_*/I*_bg,*t*_. Next, the pre-bleach intensity *Ĩ*_pre_ is the average normalized intensity in the ROI before bleaching. We define the FRAP signal as

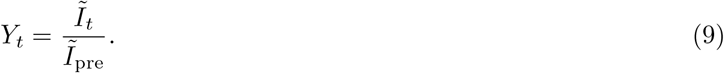

Note that the FRAP signal at *Y*_*t*=0_ at the first frame recorded after bleaching is not zero. This is for two reasons: incomplete bleaching and the inevitable delay between the first frame and the end of bleaching. During this 1.5s delay, non-bleached cytosolic myosin diffuses into the bleached region. Based on the FRAP curve *Y*_*t*_, the myosin signal can be divided into three fractions:

- The fully mobile/incompletely bleached fraction *Y*_0_.
- The slowly mobile fraction *Y*_*T*_ − *Y*_0_ (signal recovered by the end of observation).
- The immobile fraction 1 − *Y*_*T*_ (signal not recovered by the end of observation).

Previous work [19] used a different definition of the FRAP signal, namely

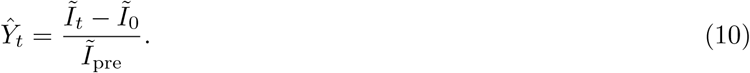

This definition leads to systematic underestimation of fluorescent recovery and overestimation of the immobile fraction, as can be seen from an example. Consider 90% effective photo-bleaching so that 10% of the pre-bleach signal is still present at the first time point post-bleach. To obtain a complete recovery using the *Ŷ*_*t*_-measure would therefore require a 10% increase in fluorescent intensity over pre-bleach levels.

### D. PRG gradient regression

In this section we give details on the model used to test for correlations between the tissue-scale myosin pattern and the pair rule genes in Fig. 1. As explained in the main text, previous work suggests that myosin is recruited to junctions between cells with differing levels of PRG expression. These cell-cell differences can be measured across the entire embryo by computing the gradient of the measured fluorescent intensity. The gradients are used as inputs in a linear regression model with the goal of predicting the myosin pattern.

Here we give a step-by-step description of this process. We enumerate the different pair rule genes (e.g. Eve, Runt and Ftz) with an index *i* = 1, 2, The level of expression of a particular gene at a position **x** on the embryo surface is a scalar function *ϕ*_*i*_(**x**). Before computing the gradients, the functions *ϕ*_*i*_ are convolved with a Gaussian kernel 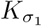 with standard deviation *σ*_1_ ≈ 1 cell diameter. This ensures that when computing the gradient, we measure differences between cells and not spurious differences between the levels within and without a cell’s nucleus. We denote the result 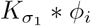. Next, we compute the magnitude of the gradient of the PRG fields, 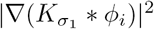. Since we consider the germ band away from the posterior pole, where the embryo is effectively cylindrical, ∇ is computed using ordinary partial derivatives. The gradient magnitude field is then smoothed again, with a second kernel of standard deviation *σ*_2_ ≈ 4 cell diameters. This is done for two reasons: First, we are interested in predicting the embryo-scale myosin distribution. Second, smoothing over small scales will reduce errors, in particular those due to imperfect alignment of data from different embryos. Smoothing makes the model more generous, as can be seen from the limit case where both the myosin and the PRG patterns are completely smoothed out - the linear regression would report perfect correlation.

The linear regression model for the myosin magnitude takes the following form:

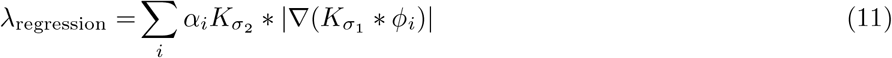

myosin magnitude is modeled by a linear superposition of PRG gradients with weights, each gene acting independently. The weights *α*_*i*_ represent the strength of each gene and are chosen by least-squares minimization. Note that the both the background levels of the PRG fluorescent signal and their image contrast are irrelevant since the former produces 0 gradient and the latter is absorbed by the fit weights. The model can be extended to allow cross terms:

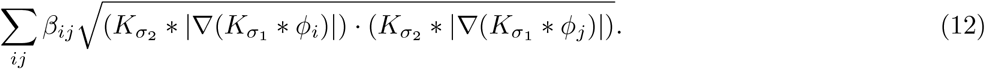

Cross terms model recruitment of myosin specifically to edges between two different PRG gene stripes. In our model, we consider three PRGs, Eve, Runt, and Ftz and allow for both linear terms and an Eve/Runt cross term.

Both the myosin and the PRG patterns depend on time *t*. This raises a potential complication, since the myosin pattern is would be expected to react to the PRGs with a delay whose duration is unknown. We carried out the linear regression to fit the myosin pattern at time *t*_myosin_ = 10 min. post VF formation when the characteristic GBE myosin pattern has already been established, but no significant deformation due to tissue flow has yet occurred. The PRG gradients are evaluated at an earlier time *t*_PRG_ < *t*_myosin_ which we can vary as an additional fit parameter. However, the PRG pattern changes little in the 20 minutes preceding *t*_myosin_ and we do not find a good agreement between the PRG-based model and the myosin distribution for any value of *t*_PRG_.

### E. Static source model

In this section we supply a quantitative model describing the behavior of junctional myosin under the hypothesis of recruitment by a static source, supporting the argument made in section 3 of the main text. During tissue flow with velocity **v**, cell junctions rotate with a rate equal to one-half of the vorticity *ω* = ∇ × **v** [5]. Note that the tissue shear does not significantly reorient cell junctions: Since GBE proceeds primarily by cell intercalation, cell shapes are not changed by the tissue-level shear.

We assume that myosin is recruited to a static source whose orientation of the source is defined geometrically: it is parallel to the direction of maximal curvature of the embryo surface. This defines the DV axis for the purpose of the measurements reported in Figs. 2-4. Myosin is assumed to be recruited to edges aligned with the source, and to detach from all edges with a constant rate 1*/τ*. These assumptions can be encoded into equations in different, equivalent ways. In main text Eq. 3 we considered a single junction, below we consider the local myosin nematic tensor constructed from detected MRJ in a small tissue patch, and in Sect. I E 1 we consider the entire angular distribution of myosin.

This yields the following equations for the local myosin tensor *m* as defined in Sect. I A 4 b:

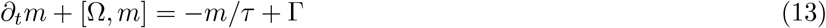

Here, Ω is the vorticity matrix Ω_*ij*_ = (*∂*_*i*_*v*_*j*_ − *∂*_*j*_*v*_*i*_)*/*2 and Γ is the source tensor, parallel to the DV axis. Advection has been neglected (see Sect. I E 2 a. In the steady state, this equation is solved by

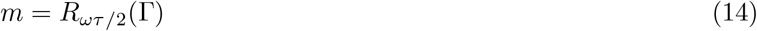

where *R*_*ωτ/*2_ represents a rotation by angle *ωτ/*2. The steady state is a good approximation for most of GBE, since after flow onset, the velocity pattern flow is relatively steady compared to the myosin lifetime *τ* ≈ 5min.

Crucially, Eq. 14 shows that the dominant eigenvector of *m*, i.e. the direction of myosin anisotropy, is given by the dominant eigenvector of Γ, rotated by an angle of *ωτ/*2. This prediction is completely independent of the magnitude of the source Γ!

#### a. FRAP-measured myosin lifetime and effective lifetime

We note that the effective lifetime *τ* in the model Eq. 13 is not necessarily equal to the time *τ*_bound_ an individual myosin motor remains bound to a junction (as measured by FRAP, for example). This can be seen by considering an example scenario in which myosin motors detach extremely rapidly, but the myosin concentration on an edge is controlled by a long-lived actor up the regulatory chain setting, for example, the rate of myosin phosphorylation. Due to the possible persistence of actors upstream of junctional myosin, the effective lifetime *τ* is larger or equal to the bound time of individual motors. This is a possible explanation for the discrepancy between the FRAP measured *τ*_bound_ and the inferred effective *τ* in main text Fig. 3.

#### 1. Model for myosin angular distribution

We next show how to predict both the mean and the standard deviation of the distribution of orientations of myosin-carrying edges. To this end, we consider a simple model for the angular distribution *m*(*t, θ*) of myosin in a tissue patch. *θ* = 0 is taken to be the local orientation of the static source. A simple model for the time evolution of this distribution, derived from the single-junction dynamics in main text Eq. 3, reads as follows:

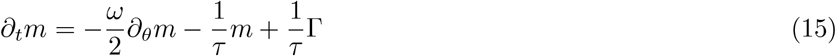

The first term describes the rotation of edges by the vorticity *ω*, the second the detachment of myosin after an effective lifetime *τ*, and the third term Γ*/τ* is the static source term, peaked around *θ* = 0. From this, one can derive Eq. 4 for the time evolution of the mean angle 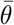. Eq. 15 is the equation used to obtain the simulated histograms in Fig. 3, with the choice 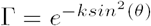. In the steady state,

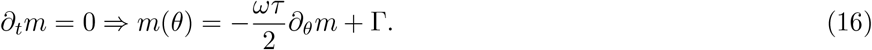

From this, the moments of *m* can be found by partial integration. The 0th moment is independent of *ω*, so that the overall amount of myosin on edges and the normalization of *m* are not affected by vorticity. For the mean *μ* and variance *σ*^2^ one finds:

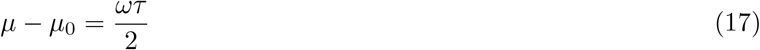

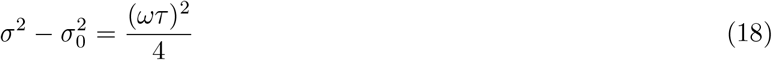

Here, _0_ denotes the values in the case of *ω* = 0. Notably, the shifts in mean and variance are completely independent of the form of the source term Γ. This means that the predictions again depend on only one parameter, *τ*.

Eq. 17 can be generalized to the case where the vorticity varies in time:

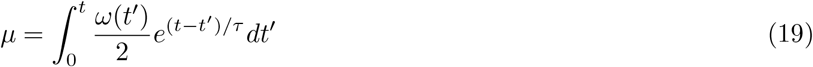

This is the equation that is used to generate the predicted myosin/DV axis angle shown in Figs. 3 and 4. One can also take into account the fact that then angle *θ* is only defined module *π* and consider moments of *e*^*i*2*θ*^ instead of *θ*, yielding similar results, for example *μ* − *μ*_0_ = tan^−1^(*ωτ*)*/*2.

##### a. Prediction of variance of myosin angular distribution

Based on our model we can predict both the mean myosin orientation as well as the variance of the angular distribution. In the main text, we presented simulated histograms (according to Eq. 15) which show striking qualitative agreement of the predicted behavior and the observed broadening of the angular distribution once vorticity sets in. In the tissue patch shown in the main text (Fig. 2), we find a change of the mean of *μ*_2_ − *μ*_1_ = 16° and a change in variances of 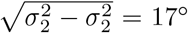 between the two timepoints analyzed, in line with the prediction 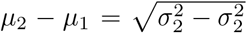 of Eq. 18. In this section, we check the prediction of Eq. 18 on the entire ensemble of *N* = 5 embryos and over the entire germ band.

However, quantitatively, the behavior of the variance is more complicated than that of the mean angle. The variance depends on the strength of the anisotropic myosin recruitment compared to the isotropic background as well as on the myosin signal-to-noise ratio, which is not the case for the mean. If the strength of the anisotropic source or the overall levels of junctional myosin (and hence the signal-to-noise ratio) decreases, as they do towards the end of GBE, the variance will increase, even in the absence of vorticity. On the other hand, a transient increase of the anisotropic recruitment leads to a decrease in variance, as is observed during the strong increase in myosin anisotropy due to strain generated by the ventral furrow during early GBE. Therefore, the vorticity effect Eq. 18 only represents one contribution to the variance. Further, the local variance depends on the size of the local tissue patch queried. If this size is chosen too large, tissue-scale gradients (e.g. of of vorticity) contribute and increase the variance. If it is chosen too small, the ensemble calculation stops making sense.

Below we show both the temporal and spatial correlation of the measured local standard deviation (in 50*μ*m × 50*μ*m windows) and the vorticity-based prediction in the germ band. We exclude a strip of 50*μ*m width around to the VF from the analysis since here the pulling effects of the VF dominate. We take the reference time for the initial variance at *t* = 10 minutes post VF initiation when the anisotropic myosin pattern is mostly established. The results of the prediction of Eq. 18 are shown in supplementary Fig. 14. The most important aspects of the spatial and temporal behavior of the variance are explained by the vorticity model.

#### 2. Additional effects in static source model

In this section, we discuss a number of effects not included in the model Eqs. 13 and 15 and why we believe they are negligible

##### a. Effect of advection

In addition to being rotated by flow, edges are also transported across the embryo surface by advection. This affects the observed spatial pattern of the myosin/DV axis angle. Advection can be accounted for by tracing the trajectory of a tissue patch back along the flow lines of the velocity field and correspond to adding a term +**v** · ∇*m* to the left hand side of Eq. 13. However, we do not do so in our vorticity-based prediction of the myosin/DV axis angle. Advection is expected to have a small effect since (a) the orientation of the geometrically defined DV axis varies very little across the embryo surface, (b) the flow is not fast compared to the myosin lifetime *τ*, and (c) around a vortex, the vorticity is approximately constant along flow lines. We can therefore neglect the effects of advection to first order and obtain a much simpler model wherein the myosin/DV axis angle at a given position is predicted from the vorticity at that same position. The residual effects of advection are one contribution to our model’s error.

##### b. Effect of myosin-feedback

If we assume that the source Γ is a function of the tension on a junction, it is likely that Γ itself could depend on the level *m* of myosin on that junction. As long as this dependence is linear, it only renormalizes the value of *τ* and has no novel effect. Here, a graphical analysis is helpful in the 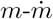 plane is helpful. Myosin detachment, i.e. 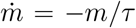 represents a straight line in this plane, whose intersections with 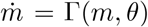 defines the steady-state of *m* on an edge of a given orientation. In the case of an *m*-independent source, this is just a horizontal line. Even if Γ depends on *m*, as long as there is only a single intersection between Γ(*m, θ*) and *m/τ*, the dynamics is qualitatively unaltered. However, the lifetime of high-myosin edges will be enhanced. Two intersections signal runaway unstable behavior in which myosin levels on an edge ratcheted up without bounds, clearly contrary to observations. In the case of three intersections, there are two stable equilibrium values and high myosin levels can sustain themselves through positive feedback. Since now myosin levels need not decay if edges rotate out of alignment with the static source, one would not expect to observe the recovery effect shown in Fig. 3.

##### c. Effects of modification of myosin lifetime by static source

The equilibrium myosin concentration 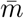 on an edge is determined by the balance of the attachment and detachment rates *k*_on_ and *k*_off_ (equivalent to the lifetime and source in the previous section): 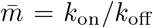. Therefore, it is possible to control the myosin distribution by either parameter, and a purported static source could influence either *k*_on_ or *k*_off_.

However, the rate by which the myosin concentration converges to the equilibrium value differs between the two scenarios. Consider an edge which rotates from an initial orientation *θ*_1_ with equilibrium value 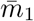 to an orientation *θ*_2_ with equilibrium value 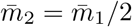. This can either happen if *k*_on, 1_ decreases by a factor of 2, or if *k*_off, 1_ increases by a factor of 2. In the the *k*_on_-case, the edge maintains an elevated myosin level for a time 1*/k*_off, 1_, but in the *k*_off_-case only for a time 1*/*(2*k*_off, 1_). This means that in the *k*_off_-case, the edge rapidly converges more rapidly back to its equilibrium value.

In order to account for the strong anisotropy of myosin we observe, with myosin on AP-edges barely above cytosolic levels, purely by a dependence of *k*_off_ on the junction orientation *θ, k*_off_ would have to be very large on junctions disaligned with the DV axis. This means that rotating junctions would rapidly lose their myosin as explained above, leading to no significant shift in the width and mean of the myosin angular distribution. Simulations similar to those shown in Fig. 3 confirm this argument.

#### 3. Principal axis of embryo-scale tension and turgor pressure

As mentioned in the discussion section, one possible candidate for the statically oriented myosin source is epithelial tension which myosin dynamics is known to be sensitive to [15]. The direction of tension agrees with the direction of the inferred myosin source. Indeed, epithelial tension in the germ band is known to be strongly anisotropic from laser ablation experiments, with higher tension on junctions parallel to the DV axis [19]. Ref. [32], using an image-based force inference algorithm, confirmed that on the scale of the entire embryo, the epithelial tension aligns with the geometric DV axis, even after the onset of tissue flow. Ref. [32] also found that the distribution of junctional myosin closely matched the epithelial tension. Strikingly, most junctional myosin is balanced: ∼80% of junctional myosin is involved in static force balance, i.e. it creates a net-zero local force.

Previous work cell-scale literature, e.g. [19], presented the anisotropic tension as a consequence of the anisotropy of junctional myosin. However, to set a static myosin orientation, the epithelial tension cannot be a pure readout of the current myosin distribution. One potential static contribution to tension anisotropy is the turgor pressure difference between the yolk within the blastoderm and the perivitlline space outside of it (visible for example during dorsal closure [33]). This normal pressure is balanced by epithelial surface stress, much like the excess pressure in an inflated balloon. Pressure, stress and geometry are linked by the Young-Laplace law. Due to the embryo’s cylinder-like geometry, the resulting surface stress is anisotropic: in a pressurized cylinder with closed ends, the stress along the azimuthal axis is twice the stress along the height axis of the cylinder [35]. Fairly generally, excess internal pressure leads to ansitropic stress parallel to principal axes of curvature [35, 52]. Interestingly, blastodermic turgor pressure leading to cortical tension has already been shown to play a crucial role in mouse blastocyst development [29]

Finally, theoretical work, Ref. [32] (recently experimentally validated in Ref. [26]) has shown how epithelial tissue can support static tension even during viscous flow. Strain-rate based recruitment can drive junctional myosin to a balanced state (with zero net local forces), such as that required to balance turgor pressure.

### F. Additional data on mutants

Fig. 15 shows a kymograph of a contracting junction in WT and *eve*, illustrating that in *eve* mutants, myosin is still associated with junction contraction. Fig. 16 shows that the myosin distribution in *eve* is significantly less anisotropic than in WT, even if the anisotropy remains clear. To test that this difference is not due to better visibility of edges in *eve*^*R*13^ due to myosin being visualized using Myosin::mCherry instead of a Myosin::GFP, we verified that this difference between WT and *eve* persists if fewer and fewer *eve* junctions are included (filtering by myosin intensity, excluding up to 3/4 detected junctions).

**FIG. 14.**
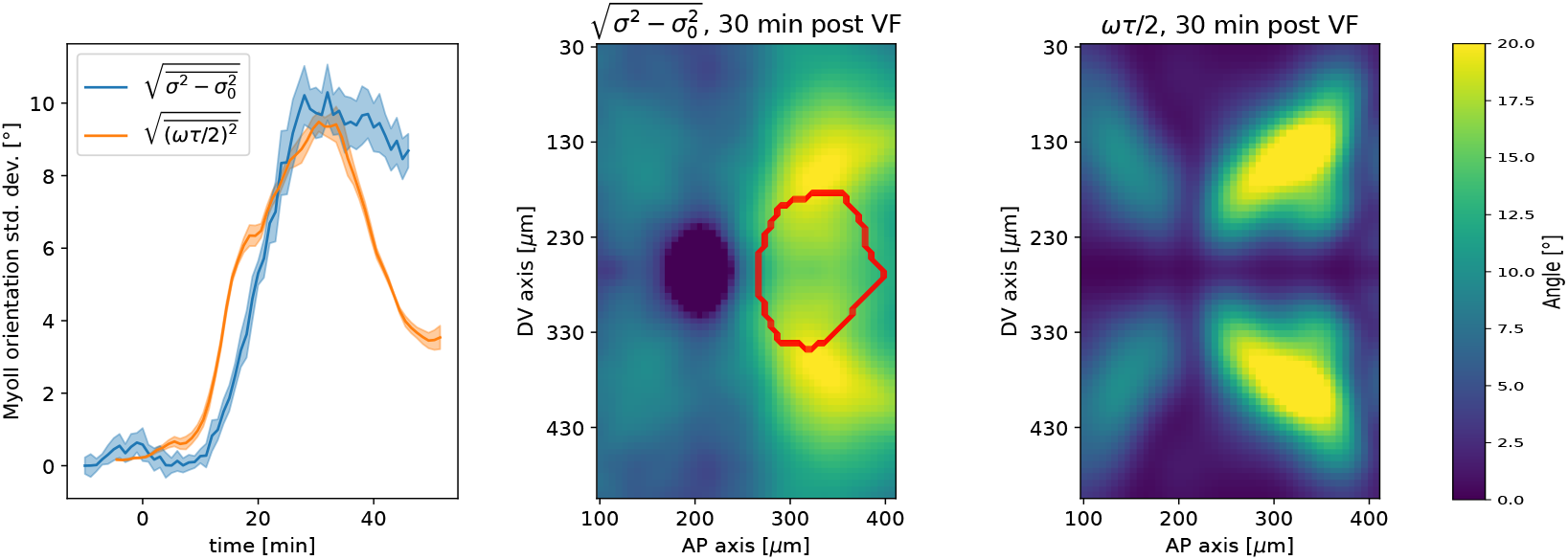
Embryo-scale prediction of myosin angular distribution width. From left to right: Spatial average over the germ band of standard deviation of myosin angular distribution and vorticity contribution to standard deviation – Heatmap of change in standard deviation of myosin myosin angular distribution, showing germ band only. Red outline indicates region of the invaginating posterior midgut – Heat map of vorticity contribution to myosin standard deviation.

**FIG. 15.**
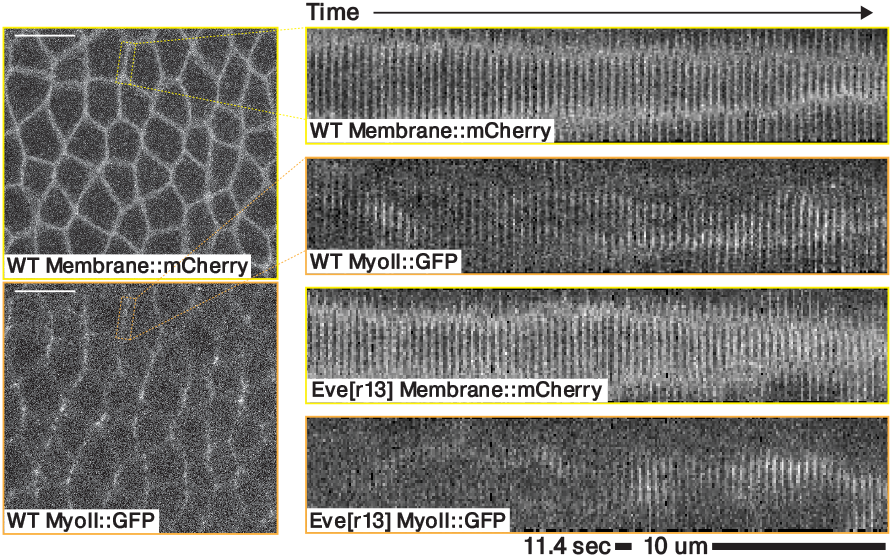
Kymograph of a contracting junction in a representative WT and a representative *eve* embryo. Both kymographs shows a junction in the germ band ∼10-20 minutes post VF initiations, marked with both a membrane and a myosin fluorescent tag.

**FIG. 16.**
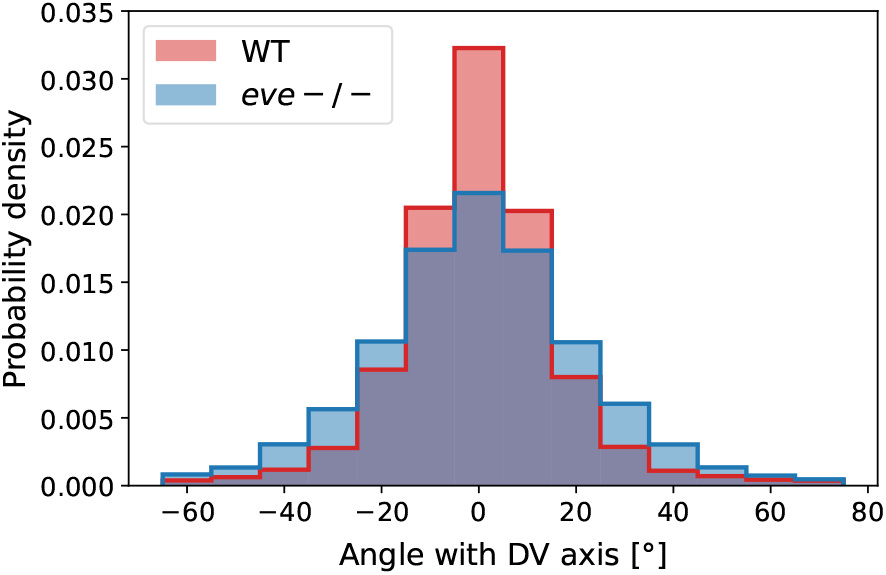
Histogram of myosin orientations in the germ band of *N* = 2 Myosin::GFP WT *N* = 2 Myosin:mCherry *eve* embryos. Data corresponds to 15 minutes post VF initiations. For each embryo, more than 7000 edges are detected. The 2-sided KS statistic (maximal area difference between cumulative distribution functions) is 0.076.

**FIG. 17.**
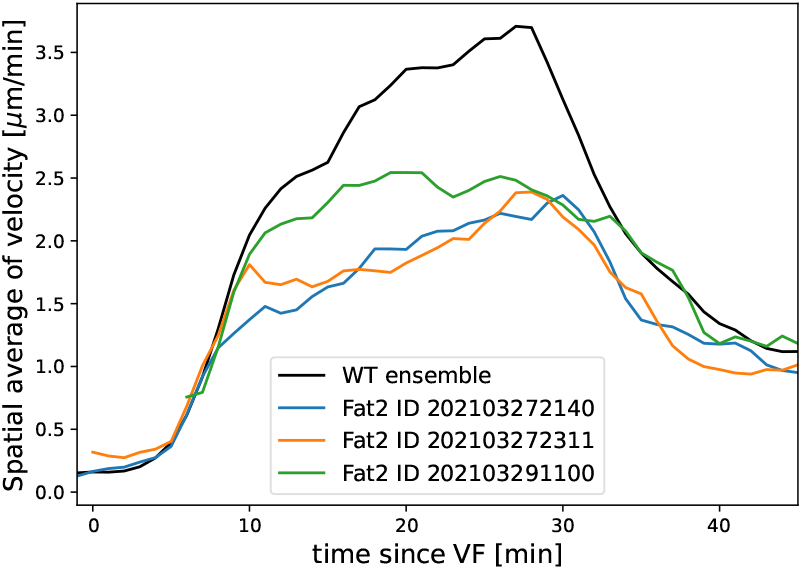
Average tissue flow velocity in WT and 3 *Fat2-RNAi* embryos. Flow in round embryos is noticeably reduced. All measurements computed using the induced metric to correct for any distortions of the cylindrical projections.

**FIG. 18.**
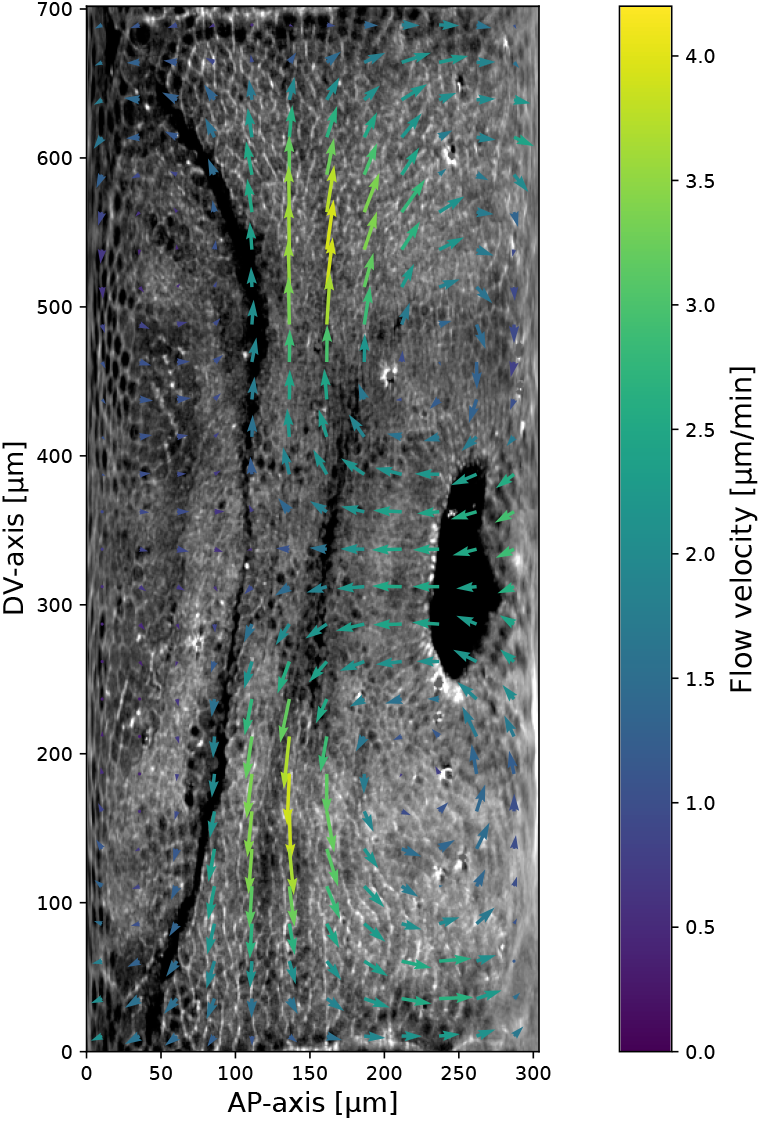
myosin and PIV field on a representative *Fat2-RNAi* embryo. Time 10 minutes post VF initiation. The signal shown is the raw myosin signal, not subjected to the cytosolic normalization procedure, to show the embryo anatomy. The PIV vortices (regions of maximal vorticity) are removed from the regions with significant junctional myosin accumulation.

## Notes

### Competing Interest Statement

The authors have declared no competing interest.

